# Integrating central and peripheral neurons in elongating multi-lineage-organized gastruloids

**DOI:** 10.1101/2020.12.29.424774

**Authors:** Zachary T. Olmsted, Janet L. Paluh

## Abstract

Human stem cell technologies including self-assembling 3D tissue models provide unprecedented access to early neurodevelopment and enable fundamental insights into neuropathologies. Gastruloid models have yet to be used to investigate the developing nervous system. Here we generate elongating multi-lineage-organized (EMLO) gastruloids with trunk identity that co-develop central and peripheral nervous system (CNS, PNS) correlates. We track migrating neural crest cells that differentiate to form peripheral neurons integrated with an upstream spinal cord region. This follows initial EMLO polarization events, and is coordinated with primitive gut tube elongation and cardiomyocyte differentiation. By immunofluorescence of multi-lineage and functional biomarkers, we evaluate EMLOs over a twenty-two day period, and apply them to investigate the impact of mu opioid receptor modulation on neuronal activity. This comprehensive study demonstrates a novel combined CNS-PNS model of early organogenesis and integration events in the trunk to benefit human biomedical research.

## INTRODUCTION

Three-dimensional *in vitro* tissue models called organoids have revealed the astonishing innate capacity of stem cells to self-organize into complex cytoarchitectures resembling *in vivo* states^1^. Directed single-organoid models of anatomical endpoints of interest, such as regions of the brain, spinal cord, gastrointestinal tract, and heart, are rapidly becoming indispensable to biomedical research for modeling physiology, development, disease, aging, and toxicity^2^. Beyond organogenesis, the study of multi-tissue interactions requires the ability to better recapitulate the morphogenetic complexity of embryogenesis. As such, a shift towards new organoid models, deemed gastruloids, that more accurately reflect early development in a multilineage, embryo-like context has emerged^3^. Pioneered using mouse embryonic stem cells (mESCs)^4^, and in one study using human embryonic stem cells (hESCs) for anterior-posterior organization^5^, gastruloids have been applied *in vitro* to demonstrate symmetry-breaking and axial elongation events^4,6^, somitogenesis^7,8^, and cardiogenesis^9^. Yet, while gastruloids provide a means to interrogate developmental processes with unprecedented detail *in vitro*, no study using mESCs or human stem cells have investigated the co-emergence of central and peripheral nervous system correlates. Such a system would constitute a major advance towards investigating CNS-PNS co-development with multiple organ precursors, and ultimately for human studies of end organ innervation.

Here we describe the first CNS-PNS neurodevelopmental model as elongating multi-lineage-organized (EMLO) gastruloids using human iPSCs. EMLO gastruloids mature from small starting aggregates, and achieve increased morphogenetic complexity and multi-system physiological relevance versus existing organoid models. We use ethnically-diverse hiPSC (ED-hiPSC) lines reprogrammed from African American, Hispanic-Latino, and Asian donors^10,11^ to demonstrate highly reproducible EMLO generation of integrated embryonic tissue precursors. In particular, we distinguish CNS and PNS correlates by spinal cord neuronal subtypes and neural crest cell (NCC) derivatives, respectively, which form in concert with a self-organizing primitive gut tube. As highly compartmentalized 3D aggregates formed without embedding, EMLOs consist of a posterior compartment with spinal cord dorsal-intermediate neuronal subtype identity by default, and an elongating anterior compartment arising from endoderm and mesoderm throughout which NCCs migrate. The primitive gut tube spans the two EMLO compartments, and acts as a hub for integration and connectivity of the spinal cord region with peripheral neurons formed by differentiating NCCs. In addition, the gut tube interacts with surrounding mesenchyme to create a permissive microenvironment for cardiogenesis that is reminiscent of *in vivo* events^12^. We characterize EMLOs over a twenty-two day period using extensive biomarker analysis to validate lineage identity and multi-cellular diversity. By small molecule interference with germ layer specification, we disrupt primitive gut tube formation and elongation, and subsequent CNS-PNS neuronal patterning. As well, we control for NCC migration from the spinal cord region by comparing full EMLOs to variants lacking the spinal cord compartment. Finally, we apply EMLOs to studies of mu opioid receptor (MOR) modulation of neuronal activity in GABAergic interneurons versus spinal sensory and spinal motor neurons, thereby demonstrating the future utility of EMLO models in pharmacological studies with agents impacting multiple organ systems. Our unique EMLO approach with human stem cells is an important step towards heterogenous 3D culture platforms with increased complexity and reproducibility that better reflects closely linked human developmental processes *in vitro*, and which is likely to benefit organ innervation studies.

## RESULTS

### Reproducible generation of elongating multi-lineage-organized (EMLO) gastruloids from African American, Hispanic-Latino and Asian hiPSC lines

To increase stem cell studies that reflect ethnic diversity (ED) in the U.S., we analyzed EMLO gastruloids using previously derived hiPSC lines from our laboratory^10,11,13–15^. Nine ED-hiPSC lines derived from the fibroblasts of three donors of self-designated African American (F3.5.2, F3.2.2, F3.3.1), Hispanic-Latino (H3.3.1, H3.1.1, H3.4.1), and Asian (A2.1.1, A2.2.1, A2.2.2) ethnicities were compared (**Supplementary Fig. 1, Supplementary Table 1**). EMLOs were generated by modifying a dual-fated neuromuscular organoid protocol (**Supplementary Fig. 1c-f**)^16^ to include mesendoderm fate by altering key initial physicochemical inductive and temporal cues (**Fig. 1a**). These modifications allowed retention of primitive mesendoderm^17^ and ectodermal phenotypes during EMLO formation that has not been previously investigated. In brief, intact 2D stem cell colonies were pretreated with CHIR and FGF2 in N2B27 induction medium for two days. After pretreatment, we generated highly uniform 3D starting aggregates (<100 μm diameter, ~300-400 cells per aggregate) by dissociation and immediate transition to low adhesion shaking cultures, bypassing static 96-well plates. FGF2, HGF and IGF-1 growth factors were included in N2B27 at the time of spontaneous aggregation. Aggregates were split 1:2 after 24 h and maintained to day 4, at which point aggregates were expanded to 100 mm dishes and maintained in N2B27 alone to at least day 22. We monitored morphogenetic properties over time by phase contrast microscopy and whole-mount immunofluorescence (IF) of tissue-cleared samples (**Fig. 1b-i**).

**Fig. 1.**
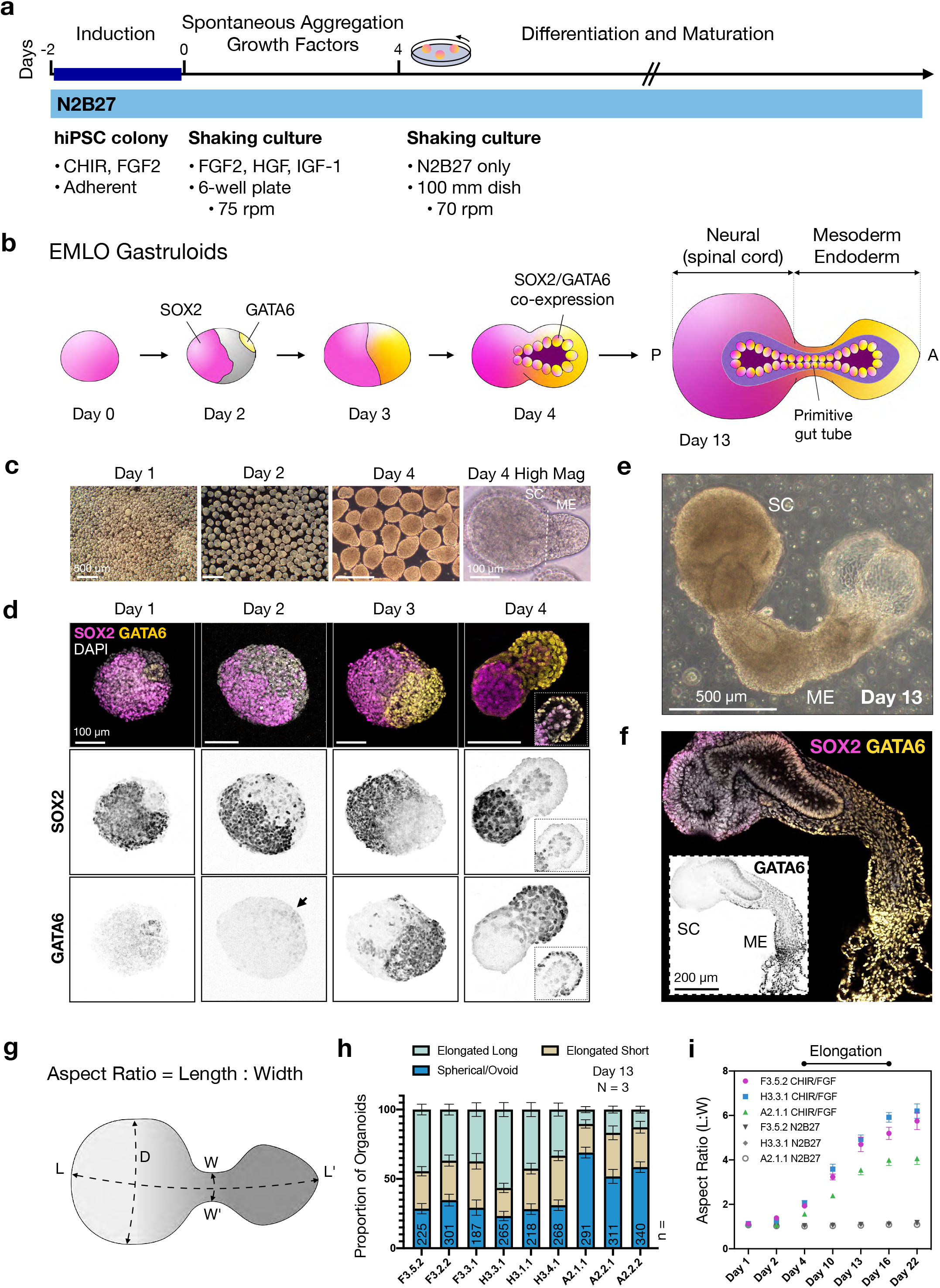
Polarized gene expression and morphology in elongating multi-lineage-organized (EMLO) gastruloids. **a**, Overview of EMLO gastruloid formation. Intact hiPSC colonies were pretreated with CHIR and FGF2 for 2 days prior to dissociation and transition to shaking culture on day 0. **b**, Schematic representation of SOX2 and GATA6 polarized expression in EMLOs over time. Early invagination of SOX2/GATA6 cells precedes formation of a primitive gut tubelike structure. **c**, Phase contrast of early EMLO shaking cultures at days 1, 2 and 4. Early compartmentalization of spinal cord (SC) neural versus mesoderm-endoderm protrusion (ME) can be visualized. **d**, SOX2/GATA6 immunofluorescence (IF) of early EMLO aggregates. Day 4 inset depicts SOX2/GATA6 early tube formation (Z-slice). **e**, Phase contrast of day 13 EMLO. **f**, SOX2/GATA6 IF in day 13 EMLOs. SOX2/GATA6 co-localization persists in the primitive gut tube. Inset is GATA6 inverted LUT. **g**, Schematic representation of the EMLO size parameters. Length (L), width (W) and SC diameter (D) were measured. **h**, Histogram of proportion of aggregates that are elongated long (L:W ratio > 3.5), elongated short (2 < L:W < 3.5), or spherical/ovoid (L:W < 2) at day 13 using nine ED-hiPSC lines. Plot corresponds to elongation efficiency. EMLO formation using all lines was performed N = 3. n = number of aggregates measured is shown for each line. **i**, Aspect ratio over time in representative ED-hiPSC lines. Elongation after pretreatment (n = 153 per line) was compared to elongation in basal medium N2B27 (n = 138 per line). Individual scale bars provided. Data reported as (mean ±s.e.m.).

SOX2 and GATA6 are two essential transcription factors involved in myriad early developmental polarization events, wherein the individual deletion of each gene is embryonic lethal^18,19^. In EMLOs, we observed polarized gene expression of SOX2 versus GATA6 as soon as 24 h after aggregation (**Fig. 1d**). A prominent GATA6 domain emerged by day 3, opposite the SOX2 domain, with some interpenetration. Notably, by day 4, we observed asymmetric bipolar aggregates that exhibited single bilaminar protrusions extending out of the SOX2 domain. While the outer cell layer was predominately GATA6+, the inner invaginated layer coexpressed GATA6 and SOX2. This mixed factor protrusion continued to elongate between days 4 and 16 (**Fig. 1e**, day 13), a critical EMLO elongation period, and ultimately produced a laminated tube-like structure that retained SOX2/GATA6 double-positivity (**Fig. 1f**). Given the similarity to embryonic structures, we refer to the SOX2 domain as the neural, or spinal cord (SC) compartment that is posterior, and the GATA6 domain as the mesoderm and endoderm compartment (mesoderm-endoderm, ME) that is anterior. We use the terminology “domain” when referring to biomarker, and “compartment” when referring to the tissue structure as a whole. To exclude any contribution of aggregate fusion to EMLO gross morphological complexity, we tracked individual aggregates and monitored elongation (**Supplementary Fig. 1g**). By this approach, we observed the emergence of a dilated vesicular structure in the anterior EMLO pole of ME that extended in length and in size to day 13 (**Fig. 1e**), after which this region resembled amorphous sacs lined by GATA6+ cells (**Supplementary Fig. 1h**).

To evaluate reproducibility of EMLOs, we measured gross size parameters and assessed formation efficiency using the morphologically distinct hourglass shape in all nine ED-hiPSC lines (**Fig. 1g-i**). The major axis length (L) was measured between the most distant points of the posterior and anterior poles, while the minor axis after elongation was defined as the width (W) of the ME compartment near the SC exit point (**Fig. 1g**). We categorized day 13 EMLOs as elongated long, elongated short, or spherical/ovoid based on aspect ratio in N = 3 repeated formation experiments (**Fig. 1h**). Replicate ED-hiPSC lines had more similar elongation efficiencies, with greater differences seen between cohorts. Amongst all lines, H3.3.1 had the highest efficiency of elongation (52.8% elongated long, 20.8% elongated short, 73.6% total). Given similar EMLO formation efficiencies within replicate lines, we chose one line from each cohort to investigate further that were F3.5.2, H3.3.1 and A2.1.1. Representative lines were previously extensively characterized genomically, transcriptomically, functionally and by protein biomarker in multi-lineage differentiation platforms^10,11^. All three lines form teratomas in nude mice^10^. Aspect ratios in EMLOs were measured over time and compared to aggregates formed without CHIR/FGF2 pretreatment (N2B27 pretreatment only) (**Fig. 1i**). Elongation was not observed in cerebral organoids generated from anterior NSCs, or in suspension neurospheres that used caudal spinal cord NSCs as starting material (**Supplementary Fig. 2**). As well, EMLO elongation was prevented by addition of 1 μM retinoic acid starting on day 2, presumably due to premature conversion to neuroectoderm (**Supplementary Fig. 2a, c**). Culture conditions that promote or inhibit EMLO elongation and efficiency are summarized (**Supplementary Table 2**).

### Self-organization of a primitive gut tube-like structure follows initial polarization in EMLOs

While a three day pretreatment of hiPSCs seeded as single cells yields homogenous adherent cultures of SOX2/Bra neuromesodermal progenitors (NMps) (**Supplementary Fig. 1c-f**),^16^ our modified protocol yielded a less restricted mesendoderm-like cellular starting material characterized by expression of SOX2, Bra, and the definitive endoderm marker FOXA2/HNF-3β (**Fig. 2a-b**). By IF of day 3 early EMLOs (**Fig. 2c**), we observed compartmentalized expression of transcription factors that orchestrate multi-lineage gene regulatory networks in development. Similarly to gastruloids^5,6^, EMLO morphological asymmetry and elongation was preceded by asymmetric polarization of cellular pools within aggregates. Cells expressing definitive endoderm markers SOX17 and FOXA2 were detected within the SOX2 domain. Bra+ cells that precede mesoderm were localized to the tip of the posterior pole by day 3. Expression of CDX2, master upstream regulator of caudal *Hox* genes, was also restricted to the SOX2 domain, opposite GATA6. Together, these data suggest that multiple germ layers co-emerge in early EMLOs and contribute to downstream morphology and reproducibility. Early cytological complexity was evident and nearly indistinguishable in all three representative lines.

**Fig. 2.**
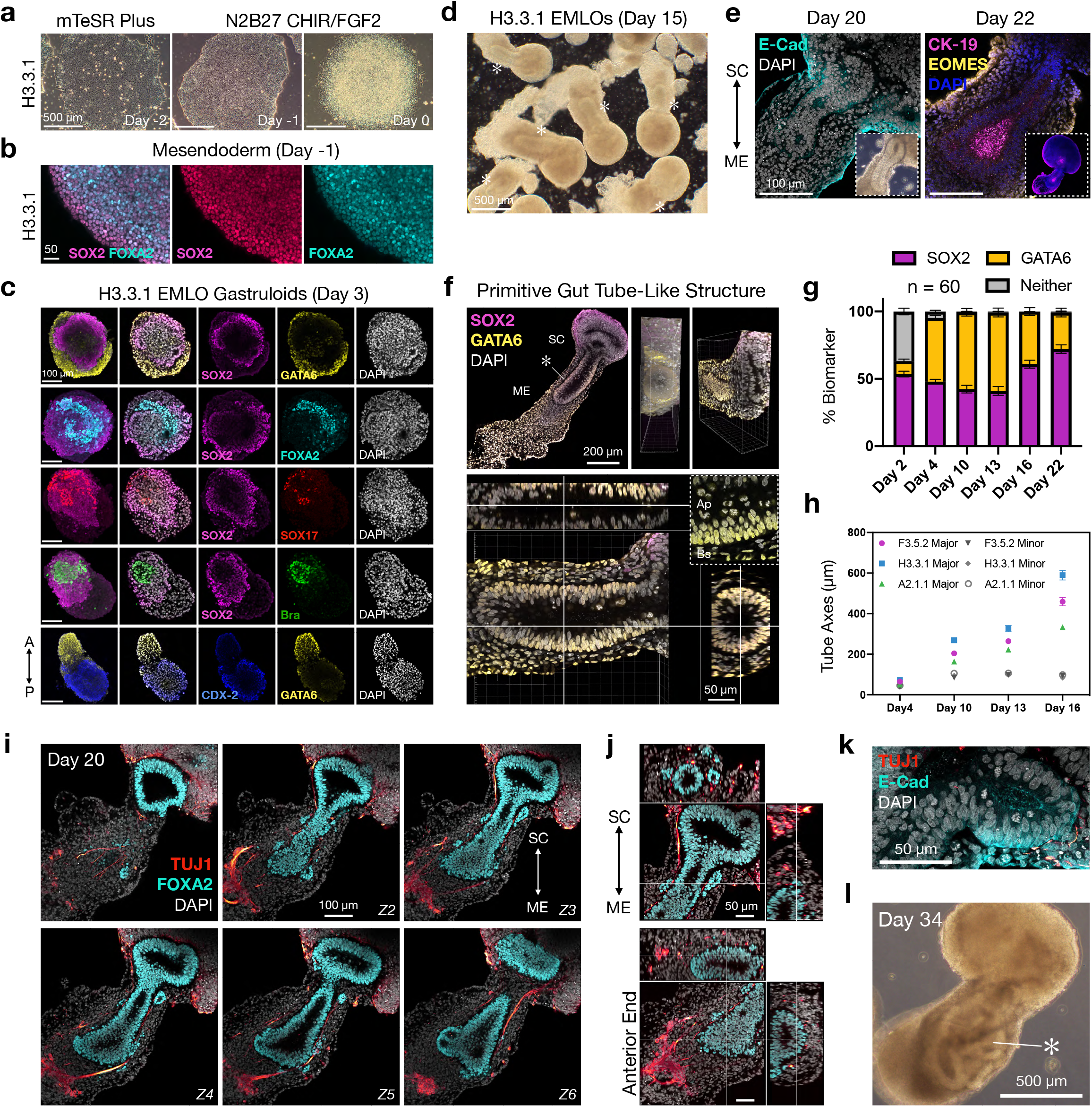
Reproducible self-organization of a primitive gut tube-like structure. **a**, H3.3.1 hiPSC colony in mTeSR Plus medium versus N2B27 + CHIR/FGF2 prior to dissociation. **b**, IF of SOX2 and mesendoderm marker FOXA2 after one day of pretreatment. **c**, Expression and compartmentalization of multi-lineage biomarkers in day 3 aggregates. SOX2, GATA6, FOXA2, SOX17, Brachyury (Bra), and CDX-2 are shown. **d**, Phase contrast of EMLO shaking cultures at day 15. Asterisk is gut tube (*). **e**, H3.3.1 phenotypic analysis of primitive gut tube-like structure at days 20 (E-Cadherin/CDH1) and 22 (Cytokeratin-19 or CK-19, and EOMES). Day 20 inset is phase contrast, day 22 inset is whole EMLO. **f**, Multi-dimensional visualization of SOX2/GATA6 primitive gut tube in day 13 EMLO. Top: confocal Z-slice. SC, ME and tube (*) are labeled (left); rotation of Z-stack 3D reconstruction (right images). Bottom: sagittal and transverse planes of gut tube. High magnification Z-slice depicts mitotic figures in apical (Ap) but not basal (Bs) epithelium. **g**, SOX2 and GATA6 distribution as percent biomarker calculated from maximally-projected Z-stacks of whole EMLOs over time. n = 60 total EMLOs measured (H3.3.1). **h**, Length of gut tube major and minor axes over elongation period in the three representative lines (n = 47 EMLOs measured per line). **i**, Z-slice series depicts robust FOXA2 expression in gut tube and associated TUJ1 neuronal fibers. **j**, Multi-dimensional visualization of FOXA2/TUJ1 at posterior (top) and anterior (bottom) aspects. **k**, High-magnification E-Cadherin/TUJ1. **l**, Day 34 EMLO phase contrast demonstrates gut tube increasing morphological complexity (*). Individual scale bars provided. Data reported as (mean ±s.e.m.).

A robust, self-organizing tube-like structure spanned the SC and ME compartments in ~100% of EMLOs that underwent elongation. This structure did not occur in the non-elongated spherical/ovoid subset. While this architectural feature was visible by phase contrast microscopy (**Fig. 2d**), we performed additional imaging studies to characterize the nature of cellular identity within the tube epithelium (**Fig. 2e, f**). E-Cadherin (CDH1) was restricted to the tube in both SC and ME compartments along with Cytokeratin-19 indicating definitive endoderm, while the surrounding tissue in ME expressed Eomesodermin (EOMES). (**Fig. 2e**). GATA6 was strictly excluded from SC, though highly expressed both in the palisading tube epithelium and the surrounding, loose mesenchyme-like tissue. These two regions in ME were separated by acellular space indicating basement membrane. This premise is supported by Z-stack 3D reconstruction, wherein the tube appears as a continuous, cylindrical structure with a central lumen containing mitotic figures at the apical aspect (**Fig. 2f**; **Supplementary Fig. 3, Supplementary Mov. 1**). Based on these data, we classified this structure as a putative primitive gut tube, hereon simply referred to as the gut tube. Gut tube identity was supported by low level SOX2 co-expression with high GATA6^4,20,21^. To roughly determine the relative proportion of neural versus non-neural lineages, we quantified the areas of SOX2 versus GATA6 domains in EMLOs over time in maximally-projected images from complete Z-stacks (**Fig. 2g**). SOX2 basal expression in the gut tube was excluded for the purpose of these measurements. We also measured the gut tube major and minor axes over time (**Fig. 2h**), which correlated with total EMLO elongation and aspect ratio (**Fig. 1i**) in support of a critical elongation period from day 4 to day 16. A FOXA2+ gut tube was evident by day 8 (**Supplementary Fig. 4a**), though became increasingly morphologically complex in close spatial relation to TUJ1 neuronal fibers (**Fig. 2i-k, Supplementary Mov. 2**) and late-stage day 34 EMLOs had meandering gut tube morphologies (**Fig. 2l**). A qualitative summary of EMLO biomarker distribution analyzed by IF is provided (**Supplementary Table 3**).

### EMLO gastruloids have default dorsal-intermediate spinal cord identity that can be ventralized

To evaluate cell and lineage identity in the EMLO neural SC compartment, we performed IF of neurons and neural crest cells, or NCCs (**Fig. 3**). By IF of β-III-tubulin (TUJ1), a neuron-specific β-tubulin isotype, we observed a dense population of neurons in SC that colocalized with TFAP2α, indicating NCC lineages and/or developing GABAergic interneuron progenitors. A small subset of neurons projected into the ME compartment, shown within GATA4+ tissue at day 13 (**Fig. 3a** top). Notably, GATA4 was excluded from the gut tube but highly expressed in the surrounding mesenchyme. At day 16, we demonstrated that the projecting neurons enveloped the GATA6+ gut tube (**Fig. 3a** bottom). A small number of TFAP2α nuclei were observed as well in close association with neuronal fibers in ME, indicating a migratory population of NCC-derived neurons or fate-biased non-neuronal NCCs that migrate along or in proximity to neuronal fibers. The number of TFAP2α nuclei in ME increased over time, while TUJ1 signal increased in both SC and ME. By day 16, TFAP2α+ neurons selforganized into a single ganglionic structure at the base of the gut tube. To validate the neural identity of SC, we showed that neural cadherin (N-Cadherin, or CDH2) is restricted to SC as opposed to the gut tube and primarily in large neural rosettes (**Fig. 3b**). Important, however, is that N-Cadherin signal was detected towards the base of the gut tube, suggesting a distal region of neural character in support of a peripheral ganglionic structure.

**Fig. 3.**
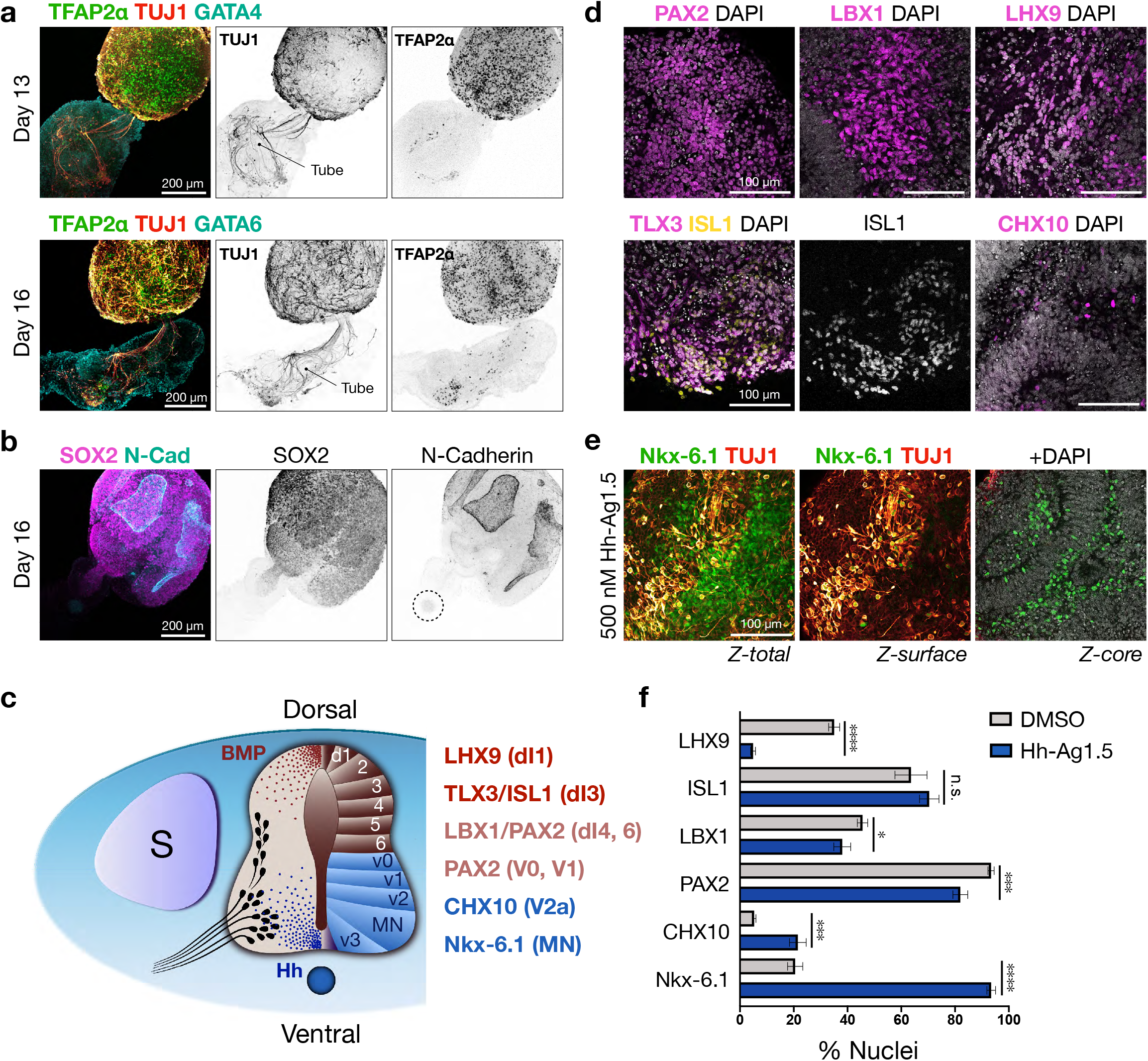
Default dorsal-intermediate spinal cord identity in EMLOs can be ventralized. **a**, IF of day 13 and day 16 H3.3.1 EMLOs depicts TFAP2α/GATA4 (top) and TFAP2α/GATA6 (bottom) counterstained with TUJ1. Position of gut tube is shown to highlight TFAP2α cell migration. **b**, N-Cadherin is restricted to SOX2 neural domain and gut tube anterior pole (dotted circle). **c**, Schematic representation of spinal cord domains along dorsal-ventral axis with ventral (Hh) and dorsal (BMP) morphogen signaling gradients. Somite (S) and notochord (ventral, blue circle) are shown. **d**, Day 22 spinal cord interneuron domain subtypes PAX2 (dI4, dI6, V0, V1), LBX1 (dI4-dI6), LHX9 (dI1), TLX3/ISL1 (dI3), CHX10 (V2a) in neural SC domain. **e**, Nkx-6.1 expression at day 22 suggests ventralization of spinal cord phenotype by addition of 500 nM Hh-Ag1.5 on day 10. **f**, Quantification of neuronal subtype biomarkers in H3.3.1 EMLOs under control (DMSO) or ventralizing (Hh-Ag1.5) conditions. LHX9 (**** p < 0.0001), ISL1 (n.s. p = 0.36), LBX1 (* p = 0.047), PAX2 (*** p = 0.0002), CHX10 (*** p = 0.0005), Nkx-6.1 (**** p < 0.0001) by unpaired t-test. n = 6,000 cells minimum counted per condition. Histogram data reported as (mean ±s.e.m.). Individual scale bars provided.

We analyzed a suite of CNS markers to further characterize the SC compartment. SC did not express anterior neuroectodermal markers OTX2 or FOXG1, presumably due to the caudal nature of the cellular starting material. However, we did detect abundant expression of several transcription factors used *in vivo* to delineate developing spinal cord progenitor and interneuron subtype domains along the dorsal-ventral axis (**Fig. 3c, f**)^22^. These were LHX9 (dI1), TLX3/ISL1 (dI3), LBX1 (dI4-dI6), PAX2 (dI4, dI6, V0, V1), CHX10 (V2a), and Nkx-6.1 (MN). By counting nuclei in H3.3.1 EMLOs, we classified the day 22 SC as having primarily dorsal-intermediate spinal cord regional identity by default due to high levels of LHX9, PAX2, and LBX1 in the context of lower CHX10 and Nkx-6.1 (**Fig. 3d-f**). PAX2 is an important marker of the intermediate spinal cord as it spans the dorsal to ventral domains. ISL1 is expressed in both dorsal dI3 interneurons and ventral spinal motor neurons, and so we used the additional marker TLX3 indicating dI3. Given the susceptibility of spinal cord regional identity to changes in opposing BMP (dorsal) and Hh (ventral) morphogenetic signaling gradients^23^, we hypothesized that addition of the potent Hh small molecule agonist Hh-Ag1.5 may shift the relative proportion of interneuron subpopulations by ventralization. Accordingly, we supplemented N2B27 with 500 nM Hh-Ag1.5 starting at day 10 and observed an significant increase in the percentage of CHX10 (*** p = 0.0005) and Nkx-6.1 (**** p < 0.0001) positive nuclei in day 22 EMLOs versus DMSO control (**Fig. 3e, f**), as well as significant decreases in PAX2 (*** p = 0.0002), LBX1 (* p = 0.047), and LHX9 (**** p < 0.0001). Together these data suggest that EMLOs have default dorsal-intermediate spinal cord identity, and that populations of neuronal progenitors and subtypes, which project from SC into ME, can be further manipulated by exogenous signaling interventions.

### CNS-PNS neuronal patterning is coupled to gut tube elongation

Given the close physical relationship between neurons and the gut tube in EMLOs, we sought to further elaborate and quantify this interaction. In parallel with an increase in the diameter of the SC compartment over time (**Fig. 4a**), we measured a corresponding increase in TUJ1 density in ME from days 13 to 22 (**Fig. 4b**). Normalized TUJ1 fluorescence was computed from complete Z-stack volumes of this isolated region as a proxy for TUJ1 density. While the gut tube was completely subsumed by TUJ1+ fibers at day 11, we observed a dense anterior bundle emanating from the base of the tube at this earlier time point prior to the formation of peripheral ganglionic structures (**Fig. 4c**). We speculate that this may be a navigational sensing mechanism to establish later stereotypic neural cytoarchitectures with high fidelity that is a common theme in development. By 3D reconstruction of high magnification Z-stacks, it is clear that neurons in ME and their projections from SC completely envelop the gut tube, but are largely excluded from the lumen and the basal lamina (**Fig. 4d**). In a subset of EMLOs, scant TUJ1 fibers were detected within the lumen. By day 16, the early formation of peripheral ganglia was observed in addition to a neuronal bottleneck effect at the transition zone between SC and ME (**Fig. 4e**). By comparison of maximally-projected Z-stacks to individual Z-slices in EMLOs at day 22, we demonstrated the retained envelopment of the gut tube and the primary large peripheral ganglion, in addition to the establishment of smaller satellite ganglia distributed throughout the ME in support of advancing neural pattern complexity (**Fig. 4f**). By 3D reconstruction of the EMLO exterior, an undulating GATA4+ ME surface interpenetrated by neuronal fibers and ganglia was evident. We refer to the anterior pole of the gut tube as the anterior intestinal portal-like region, or AIP. To determine if EMLO elongation and concomitant neuronal patterning is coordinated by the gut tube, we introduced dual SMAD inhibition by small molecules LDN 193189 (200 nM) and SB 431542 (10 μM) to inhibit mesoderm and endoderm progression during the elongation period. LDN and SB were added at day 2 in EMLO formation, a time point near the establishment of sub-aggregate polarized domains, and maintained until day 10. EMLO morphology was quantified at day 22 versus DMSO controls. At this time point, a significant proportion of EMLOs exhibited truncated ME compartments in all three lines (**Fig. 4g, h**). N = 3 repeat experiments were performed. Notably, SC cytoarchitecture that is characterized by the emergence of neurons at the base of rosettes and a bottleneck effect in the transition zone were not affected by this intervention. However, we did not detect peripheral ganglionic structures in truncated aggregates. These results suggest that ME elongation is in part driven by the gut tube and that neural patterning is related to this process.

**Fig. 4.**
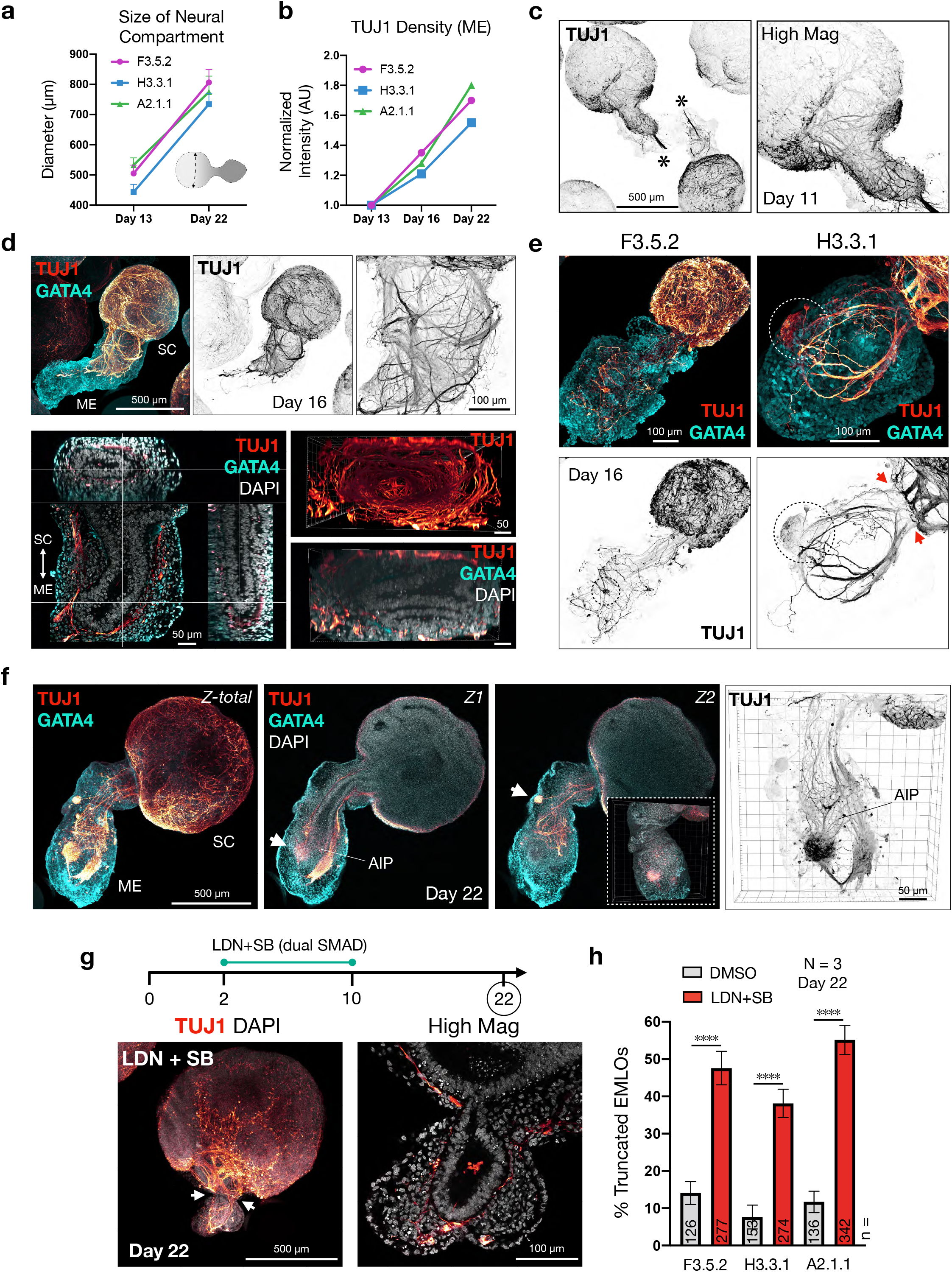
The primitive gut tube as a central organizer for EMLO neuronal patterning. **a**, Change in diameter of spinal cord (SC) compartment from days 13 to 22 (n = 47 EMLOs measured per line). **b**, Change in TUJ1 density by normalized fluorescence of Z-stack volumes between days 13, 16, and 22 (n = 3 EMLOs per time point per line). **c**, Low and high magnification of TUJ1 inverted IF at day 11 depicts neuronal envelopment of gut tube and formation of an anterior terminal bundle (*). **d**, Multi-dimensional visualization of gut tube-neuron interaction in H3.3.1 day 16 EMLO. Top: TUJ1/GATA4 (left), TUJ1 inverted high magnification (right). Bottom: sagittal and transverse planes of gut tube subsumed by neural processes (left); end-on view of neuronal tunnel (TUJ1+, top) about gut tube (DAPI, bottom). **e**, TUJ1/GATA4 IF of F3.5.2 and H3.3.1 day 16 EMLOs. Peripheral ganglion formation (dotted circle) and neuronal bottle neck (red arrows) are appreciated.**f**, TUJ1/GATA4 of H3.3.1 day 22 EMLO. Maximally-projected Z-stack (left, Z-total) is shown with two Z-slices (Z1, Z2). SC and ME labeled for orientation. Peripheral sensory ganglia form in close proximity to gut tube anterior intestinal portal-like (AIP) region. Z2 inset depicts 3D undulating GATA4 exterior with interpenetrating neurons. High magnification TUJ1 reconstruction shown (right). **g-h,** Dual SMAD small molecule inhibitors LDN 193189 (LDN) and SB 431542 (SB) before EMLO elongation phase increases truncated gut tube and ME compartment (**g**), and was quantified in (**h**) in representative ED-iPSC lines F3.5.2 (**** p < 0.0001), H3.3.1 (**** p < 0.0001), A2.1.1 (**** p < 0.0001) by unpaired t-test. n = number of EMLOs counted (day 22). Individual scale bars provided. Data reported as (mean ±s.e.m.).

### Migratory neural crest cells form PNS networks along the gut tube

Gastruloids^5,7^ and neuromuscular organoids^16^ have been shown to generate numerous cell types from more than one germ layer including NCCs. *In vivo*, competing transcriptional programs are first coactivated in NCCs, which are then lineage-biased during migration^24^. We interrogated the diversity of cell types that arise in EMLOs and their relationship to the gut tube, focusing our analysis here to NCCs^16,25^. TFAP2α and SOX10 are key markers of migratory NCCs^26^. Accordingly, In day 6 EMLOs, we observed co-expression of SOX10 and TFAP2α (**Fig. 5a**). By day 20, NCCs seemingly diverged into separate lineages during migration (**Fig. 5b, c**), consistent with *in vivo* events. We identified a small subpopulation of EMLOs lacking the SC compartment (**Fig. 5d**). In these ME-only aggregates, the gut tube was present (ISL1+), but the NCC biomarkers SOX10 and TFAP2α were distinctly lacking, consistent with NCC migration from SC into ME. To assess epithelial-mesenchymal transition (EMT), we performed IF of EMT biomarker, Vimentin, in the context of extracellular matrix components Type I and Type IV collagen (**Supplementary Fig. 5a**). EMLOs exhibited a demarcated zone of EMT at the base of SC where bottlenecking of SC neurons occurs. Type IV collagen that is an abundant component of basement membranes surrounded the basal aspect of the gut tube and lined the EMLO cell surface layer. By IF staining of PAX7 and PAX3, we validated that EMLO SC identity is largely that of dorsal-intermediate spinal cord (PAX7), while PAX3 that marks migratory NCCs originating in the SC were distributed throughout the ME (**Supplementary Fig. 5b**). As an internal control, non-elongated aggregates had dense, uniform PAX3 distribution.

**Fig. 5.**
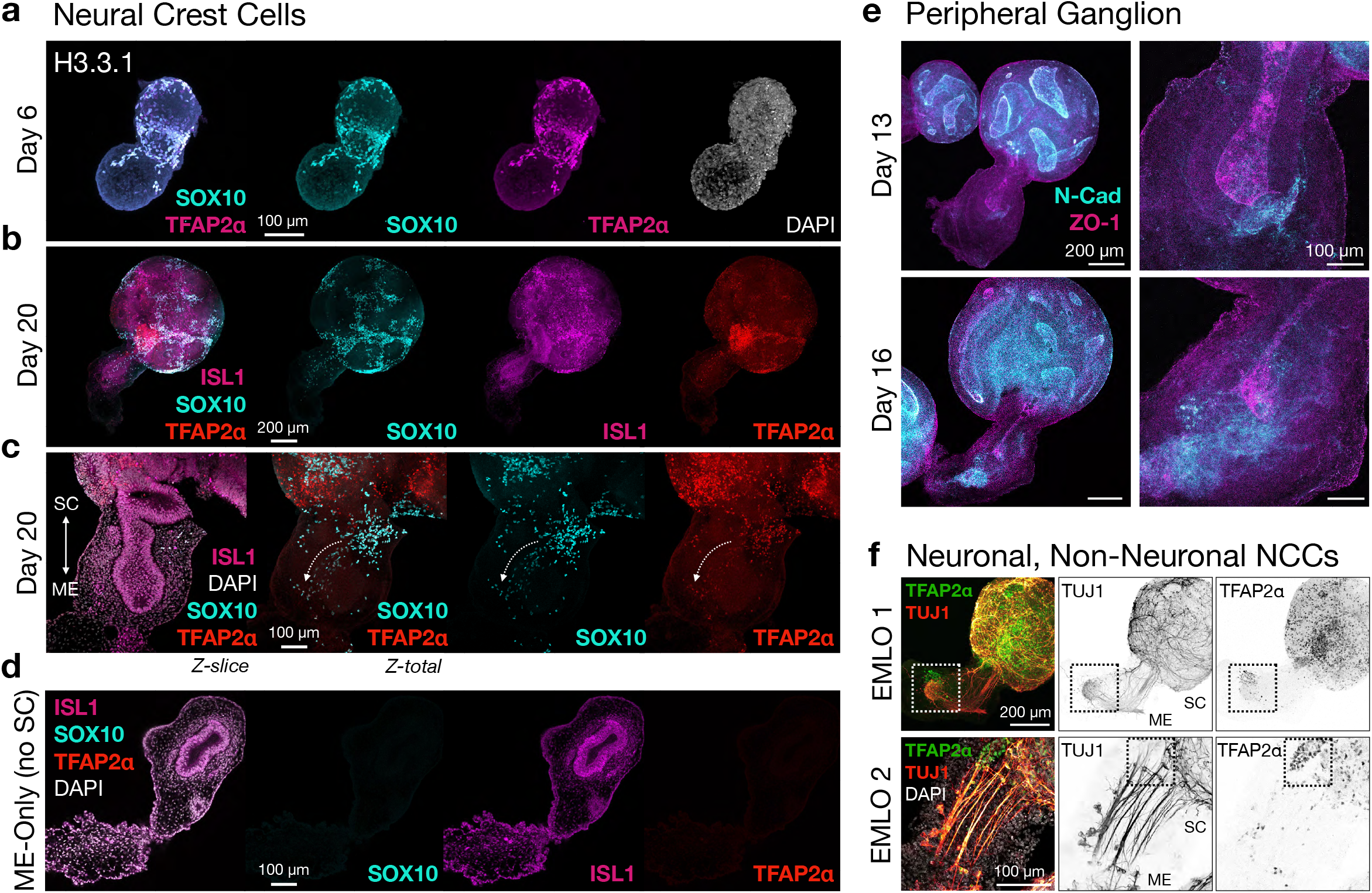
NCCs migrate to establish CNS-PNS correlates along the gut tube. **a**, TFAP2α/SOX10 double-positive cells in day 6 H3.3.1 EMLO. **b**, Diverging NCC lineages by ISL1/SOX10/TFAP2α IF in day 20 EMLO. **c**, High magnification representative day 20 EMLO. White dotted arrow depicts NCC migration over gut tube. **d**, EMLOs lacking the SC compartment do not contain NCCs, supporting NCC migration from SC into ME. **e**, N-Cadherin expression restricted to SC and the gut tube anterior pole, shown at days 13 and 16 (H3.3.1). High magnification images provided (right). **f**, Top (EMLO 1): TFAP2α/TUJ1 ganglionic colocalization at the gut tube anterior pole. A subset of TFAP2a+ cells do not express TUJ1 (dotted box), indicating non-neuronal neural crest lineages. Bottom (EMLO 2): high magnification TFAP2α/TUJ1. TFAP2α+/TUJ1-cellular pools are observed at the transition zone between SC and ME (dotted box). Individual scale bars provided.

Similarly to neurons, SOX10+ cells interpenetrated between neural rosettes and concentrated at the base of SC, with a subset migrating into ME (**Supplementary Fig. 5c**). N-Cadherin was concentrated in SC but also near the distal anterior aspect of the gut tube, and the IF signal at this region increased from day 13 to day 16 as the number of neurons populating the ganglion increased (**Fig. 5f**). The proportion of EMLOs that exhibited a prominent peripheral ganglionic structure at day 22 was quantified (**Supplementary Fig. 5d**). TFAP2α nuclei exhibited a migratory distribution in ME and concentrated at the ganglion. Two subsets of TFAP2α cells were identified. One subset co-localized with TUJ1 while the other did not. Both subsets were detected in cells emerging from SC along TUJ1 fibers as well as peripherally at the ganglion, supporting the presence of both neuronal and non-neuronal NCC derivates (**Fig. 5f**). Importantly, we detected the peripheral nervous system intermediate filament protein Peripherin in ME but not SC compartments (**Supplementary Fig. 5e**) suggesting a CNS-PNS distinction. As well, ganglionic neurons expressed ChAT that is characteristic of the enteric nervous system (**Supplementary Fig. 5f**). For all biomarkers analyzed, migratory cells were distributed along the gut tube.

### EMLO splanchnic mesoderm and gut tube interactions establish a cardiomyocyte-permissive microenvironment

The *in vivo* gut tube, and in particular the AIP, has been shown to be a putative developmental organizer of the heart^27^, likely due to reciprocal inductive signals between foregut endoderm and splanchnic mesoderm^12^. As such, we investigated the presence of the requisite tissues for cardiogenesis in EMLOs. Using IF of ISL1, a transcription factor with key roles in spinal neuron subtype diversity as well as gastrointestinal, cardiac, and neural crest development^28^, we identified several important regions (**Fig. 6a**). In day 22 EMLOs, ISL1 had low-level basal expression in the gut tube epithelium proper, but was enriched at the anterior pole and in regions of the epithelium that began to bud from the tube. Importantly, we observed a rich pool of ISL1+ cells in close proximity to the gut tube (**Fig. 6a**). The surrounding ME domain also expressed MEF2C and HAND1 (**Fig. 6b**), and TUJ1 co-localized with ISL1 pools in the primary ganglion (**Fig. 6c**). Day 13 EMLOs had some cardiac Troponin-T+ (cTnT) cells in proximity to the AIP region (**Fig. 6d**), and robust Desmin/cTnT co-expressing cardiac progenitors were present by day 20 (**Fig. 6e**), though contractile phenotypes were not investigated. Desmin+ myogenic precursors co-localized with SMI312 neuronal axons in ME (**Fig. 6f**). As well, FOXF1 that is a marker of splanchnic mesoderm was expressed predominately in cells surrounding the gut tube, and lined networks of vacant channels that coursed longitudinally along the tube towards the ganglion (**Fig. 6g**). Finally, the presence of single SOX17+ nuclei distributed along the gut tube may indicate primordial germ cell-like cells^29^(**Supplementary Fig. 4b**). Together, these findings demonstrate a diverse cellular microenvironment that is permissive to cardiac development. Multi-lineage cellular diversity and reproducible organization was observed in all three representative ED-hiPSC lines.

**Fig. 6.**
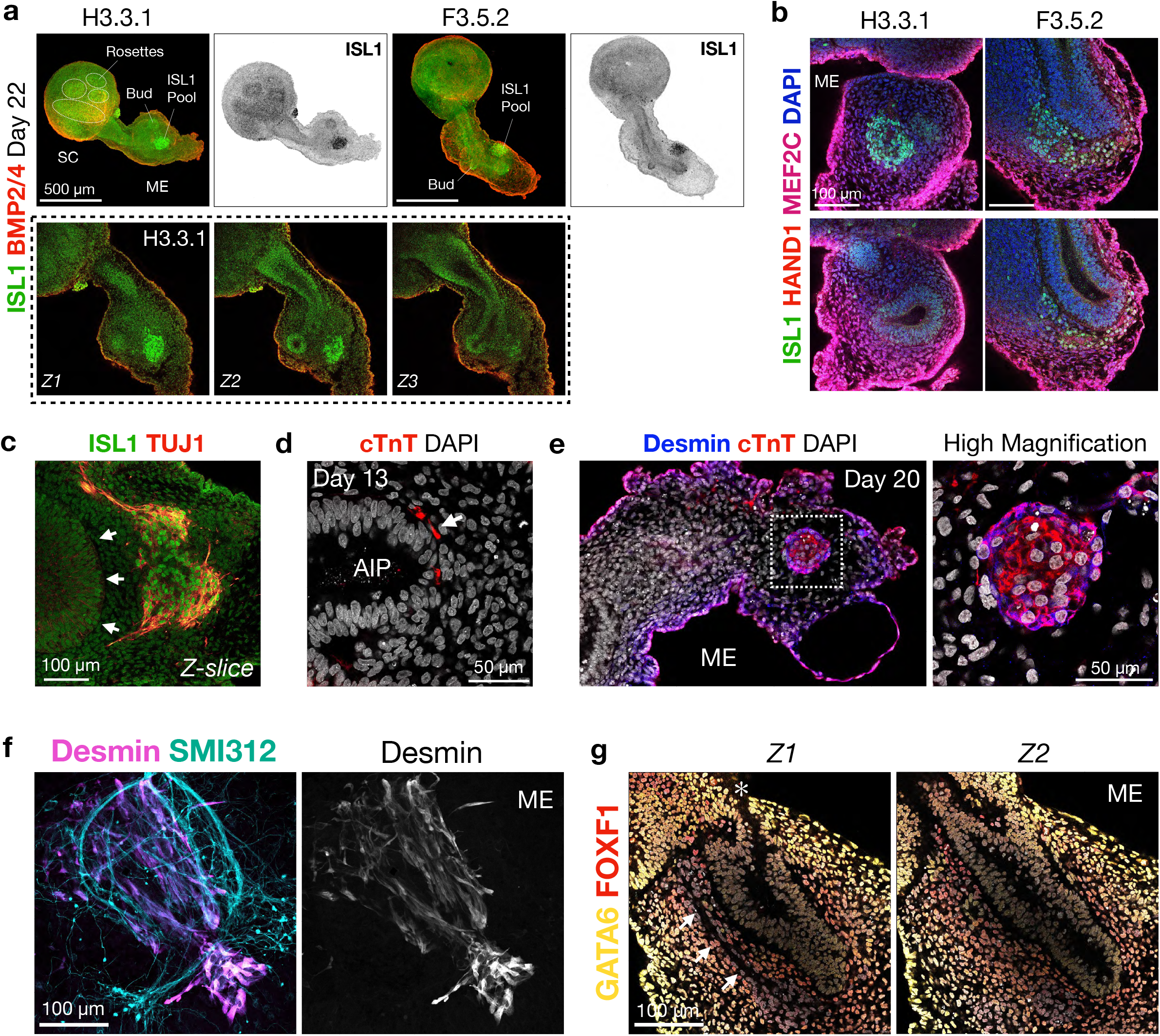
Gut tube and splanchnic mesoderm establish permissive conditions for cardiomyocyte production. **a**, Top: ISL1 and BMP2/4 IF in day 22 H3.3.1 and F3.5.2 EMLOs. ISL1+ cellular pools in proximity to the gut tube are observed in both lines, and increased expression in budding regions of the tube versus non-budding regions. Bottom: Z-slice series of H3.3.1 EMLO shown above. **b**, ISL1/HAND1/MEF2C IF in day 16 H3.3.1 and F3.5.2 EMLOs. **c**, ISL1/TUJ1 ganglionic co-localization at ISL1 peripheral pools. White arrows depict anterior gut tube. **d**, Cardiac Troponin-T (cTnT) adjacent to anterior intestinal portal-like region (day 13). **e**, Robust Desmin/cTnT expression in ME towards base of gut tube. **f**, Myogenic progenitors (Desmin) in axon-rich (SMI312) region of day 16 EMLO. **g**, GATA6/FOXF1 IF indicates splanchnic mesoderm. Asterisk marks transitory zone in cytoarchitecture (*). White arrows denote vacant channel longitudinal and adjacent to gut tube. Individual scale bars provided.

### EMLOs differentiate functional GABAergic spinal neurons that express the mu opioid receptor and have opioid-responsive firing

There are few studies with stem cells models of opioid mechanisms and virtually none that work with multiple ethnically-diverse lines. We applied EMLOs to calcium imaging studies to investigate the effect of mu opioid receptor modulation on firing activity in various spinal neuron subtypes (**Fig. 7**). In day 22 EMLOs, staining for the mu opioid receptor (MOR, or OPRM1) identified populations of OPRM1 + neurons and highlighted the apical aspect of neural rosettes (**Fig. 7a, b** top). OPRM1 signal was also distributed along the length of the gut tube lumen (**Fig. 7a** bottom). GABA synthesizing enzymes GAD65&67 were expressed in SC in congruence with dorsal-intermediate spinal cord identity and GABAergic neurotransmission that is required for the central antinociceptive effects of exogenous opioids (**Fig. 7b** bottom)^30^. We dissociated H3.3.1 EMLOs at day 22 using a neural tissue dissociation kit and formed 2D adherent neuronal networks (**Fig. 7c**). In parallel, we derived putative spinal sensory and motor neurons as positive and negative controls for MOR expression, respectively. Transcription factors BRN3A (sensory) and Nkx-6.1 (motor) were used to validate these neurons (**Fig. 7d**). 2D adherent neuronal differentiation was initiated on the same day as EMLO formation. Cultures were maintained for ten days after seeding in BrainPhys medium supplemented with BDNF, GDNF and dibutyryl-cyclic AMP. On day 32, we performed successive calcium imaging experiments using Fluo-4 AM (**Fig. 7e, f**). Baseline calcium transients were first recorded in BrainPhys medium followed by 1 μM addition of the selective MOR agonist DAMGO in the same culture. Calcium imaging was repeated, followed by addition of 10 μM of Naloxone hydrochloride to outcompete DAMGO at MOR. The percentage of neuronal somata with multiple transients above threshold over the duration of recordings (1.5 min) was quantified. In EMLO-derived cultures, the percentage of neurons with multiple transients was significantly reduced from 31 ±5% (**Supplementary Mov. 3**) to 13 ±3% (**Supplementary Mov. 4**) (mean ±s.e.m.). After Naloxone addition, the percentage of neurons increased to 30 ±5%. A similar result was observed in spinal sensory neurons, decreasing from 37 ±4% to 10 ±2%, and then increasing to 33 ±7%. Spinal motor neurons did not show a statistically significant difference between conditions, although a slight percentage increase was observed after DAMGO addition. These data support the use of EMLOs as a new early developmental model that may be informative to pharmacological studies with agents impacting many of the organs derived from the primitive structures that self-organize in the EMLO system.

**Fig. 7.**
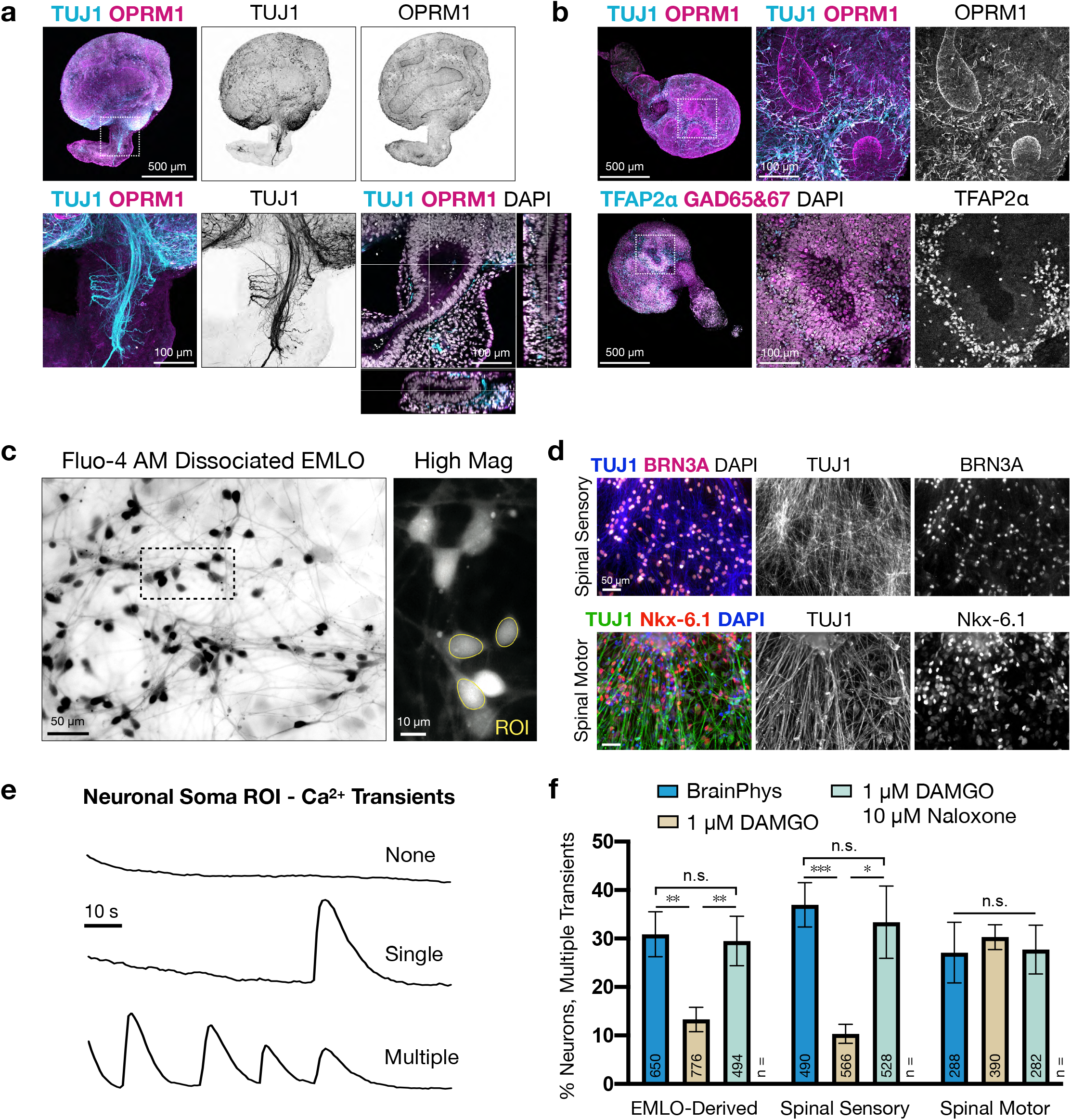
Modeling mu opioid receptor modulation in EMLO-derived neuronal cultures. **a**, TUJ1/OPRM1 IF in day 22 H3.3.1 EMLO. High magnification images provided (bottom-left) with multi-dimensional view of gut tube (bottom-right). **b**, Top: OPRM1 expression in apical aspect of neural rosettes and basal neurons. Bottom: TFAP2α expression at the basal aspect of GABAergic rosettes. **c**, Fluo-4 AM in dissociated EMLO cultures. High magnification image with example soma ROI (right). **d**, 2D adherent spinal sensory neurons (top, TUJ1/BRN3A) and spinal motor neurons (bottom, TUJ1/Nkx-6.1) used as controls for opioid-responsive firing. **e**, Examples of calcium transients quantified in dissociated EMLO neuronal soma using Fluo-4 AM. Shown are no calcium transients (none), one transient (single), or multiple transients (multiple) within a 1.5 min acquisition window. **f**, Histogram of percent of neurons with multiple calcium transients. EMLO dissociated cultures were compared to spinal sensory neurons (positive control) and spinal motor neurons (negative control). Baseline firing in BrainPhys was compared to addition of 1 μM DAMGO, followed by 10 μM Naloxone in the same cultures. n = number of neurons quantified. Individual scale bars provided. Data reported as (mean ±s.e.m.).

## DISCUSSION

In mammals, the breadth of development occurs in the post-implantation embryo, thereby limiting studies of early events that are informative for human development and disease. Gastruloids offer an ability to study the co-development of interconnected systems and organogenesis that is not currently addressable by single-organoid methods. To date, gastruloid studies have primarily applied mouse ESCs to dissect polarized developmental processes with spatiotemporal accuracy^4,6–9^. No human stem cell studies outside of this current work on EMLOs have extended early gastruloids to larger, more interconnected states composed of codeveloped integrated tissue precursors, or have evaluated the co-integration of CNS and PNS correlates in a multi-lineage study. The novel multi-system EMLO gastruloid is expected to be broadly useful for evaluating disease and toxicity impacting early CNS-PNS, gut, and heart co-developmental processes.

Perhaps the most remarkable feature that we demonstrate in EMLO gastruloids is the widespread marriage of developing CNS and PNS neurons with the self-organizing primitive gut tube (**Fig. 2 and 4**). Our data suggest a model (**Fig. 8**) in which the primitive gut tube acts as a morphological organizer along with neuronal fibers that originate in the spinal cord (SC) compartment and project into the mesoderm-endoderm (ME) compartment. A dynamic, reproducible transformation into stereotypic neuronal assemblies with peripheral ganglionic structures occurs. We espouse that neurons in EMLOs are contributed both from a posterior CNS-like region as multiple domain-specific neuronal subtypes (**Fig. 3**), and a PNS-like region that is actively patterned by the neural crest (**Fig. 5**). NMp-derived trunk NCCs *in vitro*^25,31^ have the potential to differentiate into autonomic sympathetic neurons^32,33^ and cells of the enteric nervous system^34^ that together, and in conjunction with parasympathetic and sensory neurons, comprise the PNS *in vivo*. Autonomic and enteric nervous systems are organized as diffuse networks with communicating nodes called ganglia in which neuronal cell bodies reside. In EMLOs, highly reproducible ganglionic structures emerge at the base of the gut tube and are comprised of peripheral neurons with NCC-derived expression profiles. NCCs originate in the SC compartment, become migratory, and differentiate throughout the EMLO. Similarly to *in vivo events*^24^, early NCCs co-express lineage-specific transcription factors, but commit over time to produce divergent TFAP2α, SOX10 and ISL1 NCC pools (**Fig. 5**). Migratory NCCs that populate ganglia tend to associate with preestablished neuronal fibers, and only sometimes co-localize with TUJ1 indicating both neuronal and non-neuronal subpopulations (**Fig. 5**). Peripheral neurons in EMLOs express ChAT and form in proximity to the gut tube. It is therefore possible that these structures are precursors to enteric or other peripheral plexuses. Using nine ED-iPSC lines and in-depth analysis with the three representative lines F3.5.2, H3.3.1, and A2.1.1, we consistently observed the unique CNS-PNS features present in EMLOs.

**Fig. 8.**
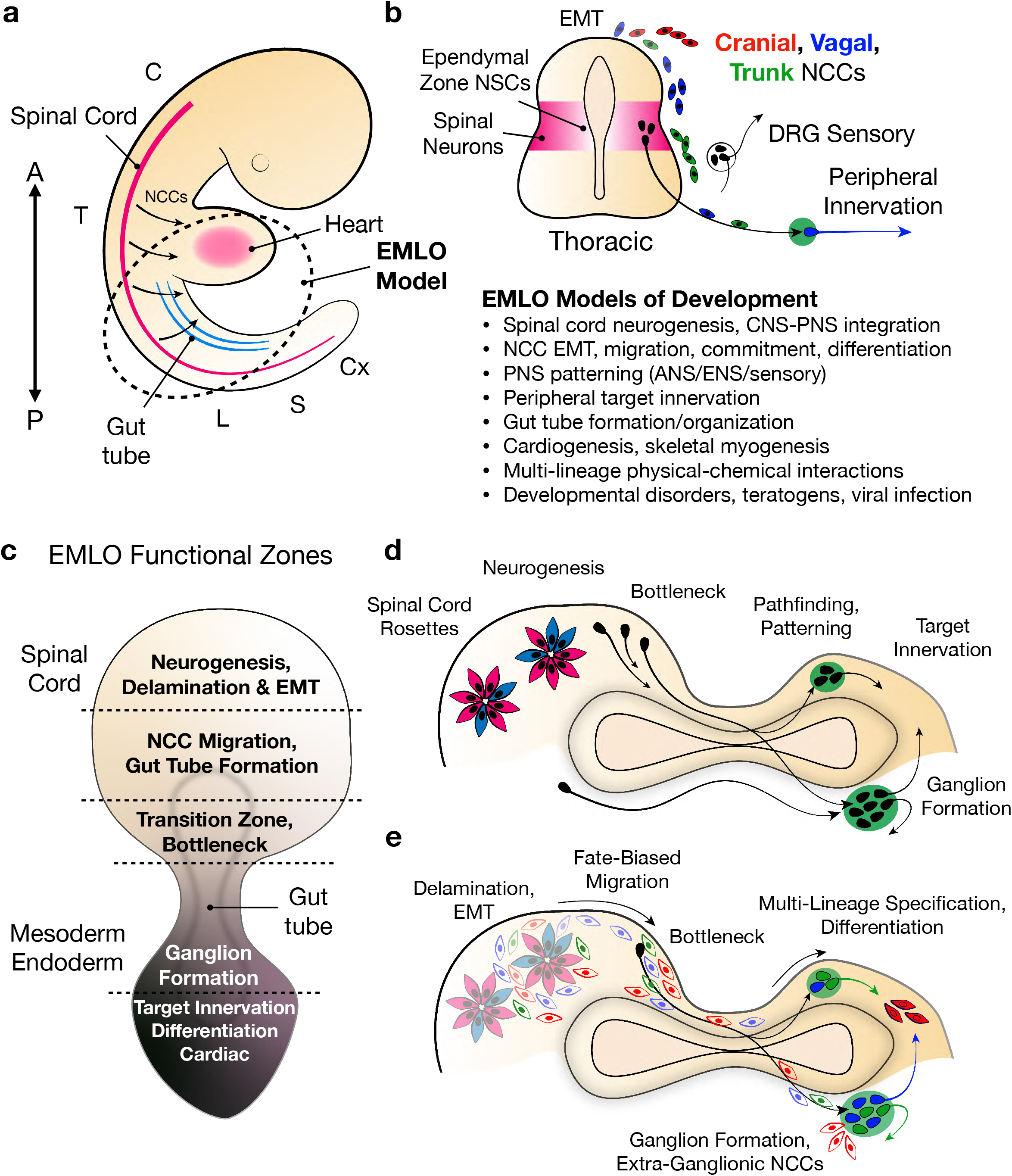
Utilization of EMLO gastruloids for human developmental studies. **a**, Schematic representation of human embryo. Cervical through coccygeal vertebral levels are labeled (C cervical, T thoracic, L lumbar, S sacral, Cx coccygeal). Primitive gut tube is shown in proximity to developing heart and spinal cord. Dotted circle represents the anatomical regions reflected in EMLO gastruloids and direction arrows are NCC migration. **b**, Neural tube cross section at thoracic level represents cranial, vagal, and trunk neural crest cell delamination, fate-biased migration, lineage commitment and differentiation. Green circle is ganglion. Below is a list of developmental processes that can be modeled in EMLOs. **c**, Segmentation model of EMLOs into approximate functional zones, although zones do overlap. **d**, Neurogenesis, migration and connectivity patterns in EMLOs including peripheral ganglion formation and target innervation. **e**, Multi-lineage neural crest cell behavior in EMLOs parallels *in vivo* events^24^.

Given the range of cell types and developmental structures observed, it is plausible that EMLOs are formed by two early pools of cellular building blocks having either mesendoderm or neuromesodermal progenitor-like identity, and whose interaction enables myriad downstream integrated tissues, including the gut tube and associated splanchnic mesoderm, that together are known to drive organogenesis^9,12,35^. Cardiac development depends on a multitude of interactions between first and second heart field progenitors and NCCs^9^, and is closely related to the foregut. *In vivo*, the anterior intestinal portal (AIP) has been shown to be a key developmental organizer of the heart by supplying inductive and mechanical signals^27^. In addition to neuronal pattern coordination, we hypothesize that the gut tube conspires with splanchnic mesoderm in EMLOs to create a microenvironment that is permissive to cardiogenesis (**Fig. 6**)^9,35^. In one previous hiPSC organoid study^36^, co-emergence of cardiac and intestinal tissue was demonstrated in spheroids, but lacked neural components and did not elongate into compartmentalized bodies or produce robust epithelial tubes or track interaction of networks over time. In Workman et al. (2016)^37^, human intestinal organoids were integrated with enteric nervous system by separately seeding with neural crest and grafting into mouse. The EMLOs generated here exhibit inherent standalone features from all three germ layers including a putative AIP, which is unique with regards to all other organoid models.

Very few studies in the literature have applied human stem cell models to opioid investigations, and virtually none that include multiple ethnically-diverse lines^38–40^. We chose to evaluate opioid biomimetic compounds in EMLOs due to the medical, financial and social burdens of diseases involving opioid use^41^. We applied EMLOs in studies examining the effect of mu opioid receptor (MOR) modulation on neuronal activity (**Fig. 7**). Dorsal lamina GABAergic interneurons play an essential role in opioid-induced central antinociception^30^, and this regional identity is present in EMLOs by default. MOR expression and responsiveness to exogenous opioids is proof-of-principle for modeling neurogenic changes of clinical and pharmacological relevance. Specifically, this analysis supports EMLOs as a first-of-a-kind model for teratogenicity studies with compounds impacting several organ systems, such as CNS, PNS, gastrointestinal, and cardiovascular, that have self-organizing precursors in EMLOs.

Together, EMLOs constitute a bridge to more systemic tissue interaction and innervation studies from gastruloids and are a valuable new approach in addition to organoids and assembloids^42^ (**Fig. 8**). The novel EMLO model enables the continued dissection of early development and, importantly, continued differentiation within an embryo-like, multi-lineage context. Although primarily tracked to twenty-two days in this study, we are continuing to track development in the EMLOs and are observing further variations off of the primitive gut. We are also exploring ability to work with the EMLOs to direct somite formation and monitoring heart field progression. This study fills an important gap in stem cell research to increase multi-ethnic representation to more accurately reflect the diverse U.S. population. As clinically-relevant human models, EMLOs are expected to offer unprecedented value in modeling organogenesis, systems interconnectivity and innervation, and the early impact of broader biomedical disease mechanisms and teratogenicity in a multi-lineage platform.

## ACKNOWLEDGEMENTS

This work was funded by SUNY Polytechnic SEED award 917035-21, and used published hiPSC lines developed through previous New York State stem cell research (NYSTEM) funding.

## AUTHOR CONTRIBUTIONS

J.P. and Z.O. conceived of the project and experimental design and analyzed data and co-wrote the manuscript. Z.O. performed EMLO formation and characterization experiments, expanded hiPSC lines for their differentiation and EMLO formation, and composed figures and illustrations.

## METHODS

### Human ethnically-diverse hiPSC lines and culture conditions

Nine ethnically-diverse (ED) hiPSC lines were previously reprogrammed from the fibroblasts of three individuals from samples banked at Coriell^10^ and were extensively characterized by our laboratory^11,13,14^. Coriell donor catalog numbers are as follows: GM22268 (F-3; African American male, foreskin, 1 day); AG08498 (A-2; Asian male, foreskin, 1 day); GM22186 (H-3; Hispanic-Latino male, foreskin, 1 day). hiPSC lines were checked for normal karyotype. Details of ED-hiPSC characterization are provided in **Supplementary Table 1**. hiPSCs were maintained in pluripotency medium mTeSR Plus (STEMCELL Technologies) supplemented with 1x penicillin-streptomycin (P-S) on hESC-qualified Matrigel (1:100 dilution; Corning) at 37°C, 5% CO_2_. Cultures were passaged 1:6 in 6-well plates every 4-7 days using Gentle Cell Dissociation Reagent and cryopreserved in mFreSR according to manufacturer’s instructions (STEMCELL Technologies). All lines were expanded and stored at low passage number.

### EMLO gastruloid formation and signal manipulation

Our gastruloid protocol begins with pretreatment of 2D adherent hiPSC colonies maintained in mTeSR Plus pluripotency medium. At ~60% confluency, mTeSR Plus was replaced with the N2B27 basal medium supplemented with 3 μM CHIR99021 (Tocris Bioscience) and 40 ng/ml basic fibroblast growth factor bFGF/FGF2 (R&D Systems). N2B27 basal medium: 1:1 DMEM/F-12:Neurobasal Plus medium, 2% (v/v) B27 Plus supplement, 1% (v/v) N2 supplement, 1x GlutaMAX, 1x MEM Non-Essential Amino Acids, 1x P-S. Colonies were maintained in pretreatment medium for two days, with fresh culture medium replenished each day. On the day of dissociation (day 0), wherein migratory cells became visible at the colony edge, cells were dissociated with 1:1 Accutase:HBSS (Ca-Mg free) at 37°C for 5 min followed by manual trituration with a P-1000 pipette. 6-well plates were pretreated with Anti-Adherence Rinsing Solution (STEMCELL Technologies) for 5 min incubation at room temperature followed by two rinses with equal volumes of fresh HBSS. Cells were suspended in N2B27 supplemented with 10 ng/ml FGF2, 2 ng/ml IGF-1, 2 ng/ml HGF (R&D Systems) and 50 μM Y-27632 (Tocris Bioscience), and single cell suspensions were distributed at 2 x 10^6^ cells/ml in 2 ml per well. Well plates were transitioned to shaking suspension culture using an orbital shaker at 75 rpm clockwise in a humidified incubator. The next day, one-half volume of medium was replaced with fresh medium N2B27 supplemented with 2 ng/ml IGF-1, 2 ng/ml HGF. On day 3, the entire volume of medium was replenished and shaking cultures were maintained to day 4. On day 4, individual wells were transferred 1:1 to 100 mm petri dishes pretreated with Anti-Adherence Rinsing Solution in fresh N2B27 only (7-8 ml per dish) and maintained at 70 rpm. One-half volume of fresh N2B27 was replenished every 3-4 days. EMLO gastruloids were maintained at least to day 22, and up to day 34.

To test the effect of dual SMAD inhibition on ME compartment morphology, 200 nM LDN 193189 and 10 μM SB 431542 (Tocris Bioscience) was added during a critical endoderm expansion/elongation period between days 2 and 10. LDN and SB were excluded after day 10 and EMLOs were quantified at day 22. Dorsal-ventral regional identity of spinal domains was manipulated by addition of 500 nM Hh-Ag1.5 (Cellagen Technology) on day 10 after elongation had occurred and maintained to day 22 for imaging and quantification by cell counting.

### Spinal sensory and motor neuron derivation

Spinal sensory neurons^43^ and spinal motor neurons^13,14^ were derived similarly to as previously described. For nociceptor sensory neurons, hiPSCs were seeded onto freshly-coated Matrigel 20,000-40,000 cells/cm^2^ in mTeSR Plus with 10 μM Y-27632. When cells were confluent (day 1 of differentiation), medium was changed to Knockout DMEM with 15% (v/v) Knockout Serum Replacement, 1x GlutaMAX, 1x MEM NonEssential Amino Acids, 1x P-S. Dual SMAD inhibition was achieved by 100 nM LDN 193189 and 10 μM SB 431542 days 0-5 and nociceptors were induced by addition of 3 μM CHIR 99021, 10 μM SU5402, 10 μM DAPT between days 2 and 10. Beginning at day 4, N2 supplement was added every other day at 25% increments to day 10. At day 20, cultures were maintained in BrainPhys supplemented with N2/B27 and 10 ng/ml BDNF, 10 ng/ml GDNF (R&D Systems), 1 μM dibutyryl-cyclic AMP (dbcAMP).

For spinal motor neurons, restricted neuromesodermal progenitors (NMps) were induced in 2D adherent hiPSC colonies (~60% confluency) by culture in N2B27 supplemented with 40 ng/ml FGF2, 40 ng/ml FGF8 (R&D Systems), 2 μM CHIR 99021, 10 μM DAPT (Tocris Bioscience), 10 μM SB 431542, 100 nM LDN 193189, 0.36 U/ml heparin^13,14^ (Millipore Sigma). Medium was changed daily for four days. At day 4, NMps were split in N2B27 supplemented with 100 nM RA, 200 nM Hh-Ag1.5 and maintained to day 20 at which point, BrainPhys supplemented medium described above was added. Spinal sensory and spinal motor neuron differentiations began the same day as EMLO formation. Day 22 cultures were passaged to chambered coverglass and cultured for ten days prior to calcium imaging.

### Phase contrast imaging, whole-mount immunofluorescence, and tissue clearing

For phase contrast microscopy, samples were imaged at room temperature directly in culture plates. Images were acquired using a Zeiss Invertoskop 40C (5x/0.12 CP-Apochromat, 10x/0.25 Ph1 A-Plan and 20x/0.30 Ph1 LD A-Plan, 40x/0.50 Ph2 LD A-Plan) mounted with an Olympus DP22 color camera and cellSens acquisition software. For whole-mount immunofluorescence preparation, EMLOs were pooled on the day of fixation, rinsed once with 1x phosphate-buffered saline (PBS) and fixed in 10% neutral buffered formalin solution at 4°C for 2 h. EMLOs were rinsed three times in 1x PBS for 5 min. Samples were then permeabilized by three successive incubations in 0.2% Triton X-100 in 1x PBS (PBST) for 20 min at 4°C, and blocked overnight in 1% bovine serum albumin (BSA) in PBST. The next day, samples were distributed to 12-well plates in 1 ml blocking solution per well. Primary antibodies were added to requisite dilutions (see Supplementary Information). Plates were left rocking at 4°C for 48-72 h, then rinsed three times in blocking solution followed by three times in PBST for 5 min at room temperature in 2 ml centrifuge tubes. Species-matched secondary antibodies were incubated 1:10,00 with NucBlue fixed cell stain (Invitrogen) directly in 2 ml centrifuge tubes, rocking overnight at 4°C. For triple-antibody stains, goat anti-mouse or goat anti-rabbit Cy5 secondary antibodies were added the next day after removal of donkey anti-goat AlexaFluor secondary antibody by additional wash steps. All excess secondary antibodies were ultimately removed by two wash steps in blocking solution followed by two wash steps in PBST. Stained and rinsed EMLO samples were equilibrated in 0.1 M phosphate buffer (PB: 0.025 M NaH_2_PO_4_, 0.075 M Na_2_HPO_4_, pH 7.4) by three successive incubations of 5 min at room temperature. EMLOs were then post-fixed in 10% neutral buffered formalin for 20 min at 4°C, and rinsed three more times in 0.2 M PB. To clear samples, PB was aspirated and replaced with 200 μl of 88% Histodenz solution (w/v) in 0.2 M PB. Samples were left in the dark at 4°C for 24-48 h and then mounted on glass slides, sealed in clear nail polish.

Samples were imaged on a Leica confocal TCS SP5 II system in conjunction with Leica Application Suite Advanced Fluorescence software. The SP5 II system was equipped with 10x/0.30 HCX PL FLUOTAR air, 20x/0.70 HC PL APO CS air or immersion, and 40x/1.25 HCX PL APO immersion objective lenses. Complete or partial Z-stacks were acquired at ~2 μm separation distance.

### Dissociation cultures and adherent neuron immunofluorescence

Day 22 EMLOs (~25 count) were pooled and dissociated using the Neural Tissue Dissociation Kit (P, palpain) according to manufacturer’s instructions (Miltenyi Biotec) and similarly to previous reports^44^. Samples were manually triturated with a P-1000 pipette. Dissociation was confirmed by visual inspection. Spinal sensory and spinal motor neurons used as controls for calcium imaging were derived as detailed above. On day 22 in differentiation, adherent cultures were detached with 1:1 Accutase in HBSS and seeded into chambered wells of Matrigel-coated coverglass. At the same time, day 22 dissociated EMLOs were seeded similarly. Parallel cultures were maintained in BrainPhys supplemented with N2, B27 and 10 ng/ml BDNF, 10 ng/ml GDNF, 1 μM dbcAMP. One-half volume of media was replenished every 3-4 days.

For immunofluorescence of adherent spinal sensory and motor neurons, day 32 adherent cultures were rinsed once in 1x PBS and fixed with 10% buffered formalin for 30 min at 37°C. Samples were rinsed again in 1x PBS, permeabilized for 5 min in 0.1% Triton X-100, and blocked for 30 min in 1% BSA fraction V (1x PBS) at room temperature. Primary antibodies were applied in 1 ml fresh blocking buffer and incubated at 4°C overnight. Samples were rinsed thoroughly in 1x PBS before applying immunoglobulin- and species-matched secondary antibodies for 1 h in the dark with NucBlue fixed cell stain (4°C). Cells were imaged directly in chambered cover glass wells. Wide field fluorescence microscopy was performed using a Zeiss Axio Observer.Z1 inverted fluorescence microscope (20x/0.8 air and 63x/1.4 oil Plan-Apochromat DIC objectives). Images were acquired using an Hamamatsu ORCA ER CCD camera and Zeiss AxiovisionRel software (ver. 4.8.2). For adherent cultures, 10-to 20-slice Z-stacks were gathered at 1 μm separation distance and compressed using the Extended Focus feature. If necessary, images were adjusted linearly for brightness in Keynote or ImageJ.

### Mu opioid receptor modulation and calcium imaging

On the day of live cell calcium imaging, 50 μg Fluo-4 AM cell-permeant dye (Invitrogen) was diluted in 10 μl of 20% Pluronic F-127 in DMSO. Fluo-4 AM was diluted 1:1,000 in BrainPhys medium and added to coverglass chamber wells after rinsing cells once in HBSS. Samples were incubated at 37°C for 30 min. After incubation, Fluo-4 AM medium was removed, washed once in HBSS and fresh BrainPhys without phenol red was added. Samples were imaged directly on the Zeiss Axio Observer.Z1 system. Time lapse series were acquired at 50 ms exposure using a 488 nm LED at 200 ms intervals for 1 to 1.5 min duration. After imaging baseline activity in BrainPhys, 1 μM DAMGO (MOR agonist, 10 mM stock in water; Abcam) was added directly to chamber wells and imaged again. The competitive opioid receptor antagonist Naloxone hydrochloride was then added at 10 μM (10 mM stock in water; Abcam). EMLO dissociated cultures, spinal sensory and spinal motor neuron samples were imaged in succession to control for time of Fluo-4 AM cell loading. Time lapse series were quantified in ImageJ.

### Quantification and statistical analysis

Raw data were compiled in Microsoft Excel (v16.16.27) and exported to GraphPad Prism (v9.0.0) for plotting and statistical analysis. Data are reported as (mean ±s.e.m.) and analyzed using unpaired two-tailed t-test. One way ANOVA was used to determine statistical significance between groups of cell lines. EMLO cell nuclei were manually counted using ImageJ to quantify immunofluorescence data using Z-slices at least 20 μm apart to avoid double counting. ****p < 0.0001, ***p < 0.001, **p < 0.01, *p < 0.05, n.s. not significant (a = 0.05). Power analysis was not performed. Detailed information for each experiment is provided in Results and Figure Legends.

### Figures

Figures for this manuscript were made in Keynote (v10.3.5) and Adobe Illustrator Creative Commons 2020. Data plots were generated using GraphPad Prism 9.

