## Supplementary Information File for "Integrating central and peripheral neurons in elongating multi-lineage-organized gastruloids"

#### SUPPLEMENTARY FIGURE LEGENDS

**Supplementary Fig. 1 Induction and tracking of pluripotent ED-iPSC representative lines to neuromuscular trunk organoids and EMLO gastruloids. a**, Phase contrast images of low passage number F3.5.2, H3.3.1, A2.1.1 pluripotent hiPSC colonies. **b**, Pluripotency markers SOX2, SSEA4 and SSEA3. **c**, Summary of neuromuscular trunk organoid protocol (Faustino Martins et al., 2020). **d**, Single cell NMp induction for three days. **e**, Rich SOX2+/Bra+ NMp population after three days of single cell CHIR/FGF2 induction in N2B27. **f**, Day 10 neuromuscular trunk organoids after transition to 60 mm dish. **g**, Representative early (top) and single-EMLO tracking (bottom) by phase contrast microscopy. Asterisk (*) indicates vesicle formation. **h**, Large dilated anterior vesicles in day 13 EMLO (left) that later appear as amorphous sacs (day 16) lined by GATA6 cells (right images). See also **Fig. 2d**. Individual scale bars provided.

**Supplementary Fig. 2 Non-elongating stem cell models. a**, Experimental overview of small molecule manipulation to prevent elongation. **b**, Phase contrast image of spinal cord NSC-derived neurospheres (NS). **c**, NMp aggregates coaxed into neurospheres by 1 μM retinoic acid (RA) added at day 2. **d**, Cerebral organoids do not elongate. Protocol adapted from Trujillo et al. (2019), with additional use of STEMCELL Technologies Cerebral Organoid kit. Individual scale bars provided.

**Supplementary Fig. 3 Annotated EMLO gastruloid.** Labeled EMLO depicting key structural and morphogenetic features. CIP, caudal intestinal portal; AIP, anterior intestinal portal. See also **Fig. 2f**. Scale bar is 50 μm.

**Supplementary Fig. 4 Definitive endoderm and neural crest cells emerge in EMLOs. a**, Primitive gut tube formation and early NCC migration in elongated day 8 EMLO. FOXA2/SOX2/TUJ1 (top), SOX10/TFAP2a/TUJ1 (middle), SOX1/ISL1/TUJ1 (bottom) Z-slices are shown with DAPI. Leftmost image is maximally-projected Z-stack (no TUJ1 or DAPI). **b**, SOX17 definitive endoderm in day 16 EMLO. Similar staining patterns are seen between elongated and spherical/ovoid aggregates. Individual SOX17+ cells are dispersed in ME along the gut tube, potentially indicating primordial germ cell-like cells. SOX17 inverted LUT shown. Individual scale bars provided.

**Supplementary Fig. 5 NCC epithelial-mesenchymal transition, migration, and ganglion formation in EMLOs. a**, Left: EMLO exterior Z-slice depicts transition zone from SC to ME marked by Vimentin indicating active EMT. Inset is high magnification. Right: EMLO interior Z-slice demonstrating basement membrane of gut tube marked by Collagen IV. Inset is high magnification. **b**, PAX7/PAX3 IF supports spinal cord identity of neural compartment and peripherally distributed PAX3+ cells in EMLOs. Non-elongated organoid shown to the right has uniform PAX3 distribution. **c**, Maximally-projected Z-stack (Z-total) of SOX10 IF highlights NCC migration into ME. Single Z-slice (right) shows interpenetration of SOX10+ cells between neural rosettes. **d**, Histogram quantification of EMLOs with neuronal ganglionic structures in ME in F3.5.2 (n = 25), H3.3.1 (n = 31), and A2.1.1 (n = 19) ED-hiPSC lines at day 22 (median). **e**, PNS marker Peripherin is expressed in neurons in ME but not SC. **f**, TUJ1/ChAT expression. ChAT is expressed in the ganglion. Maximally-projected Z-stacks are shown with Z-slices of the tube (top) and ganglion (bottom) planes. Individual scale bars provided. Data reported as (mean ± s.e.m.).

#### SUPPLEMENTARY MOVIES

**Supplementary Mov. 1** Z-stack of day 13 EMLO SOX2/GATA6, 40x magnification, 7 frames/s. See also **Fig. 2f.**

**Supplementary Mov. 2** Z-stack of day 20 EMLO TUJ1/FOXA2, 40x magnification, 7 frames/s. See also **Fig. 2i, j**.

**Supplementary Mov. 3** Fluo-4 AM in EMLO-derived adherent neuronal cultures (BrainPhys baseline). 20x magnification, 17 frames/s, 50 ms exposure at 200 ms interval acquisition (1.5 min).

**Supplementary Mov. 4** Fluo-4 AM in EMLO-derived adherent neuronal cultures (1 µM DAMGO). 20x magnification, 17 frames/s, 50 ms exposure at 200 ms interval acquisition (1.5 min).

#### SUPPLEMENTARY TABLES

**Supplementary Table 1. Ethnically-Diverse hiPSC Lines**

| **Line** | **Passage number** | **Pluripotency** | **Teratoma** | **Multi-lineage differentiation** | **hiPSC bulk RNA-Seq** | **ChIP-Seq** | **EMLO form.** |
|---|---|---|---|---|---|---|---|
| **F3.5.2** | 15-25 | IF, RT-PCR | Yes | Tomov et al. 2016 | Tomov et al. | Tomov et al. | Yes |
| F3.2.2 | 7-8 | IF, RT-PCR | Not tested | This study | Not tested | Not tested | Yes |
| F3.3.1 | 10-11 | IF, RT-PCR | Not tested | This study | Not tested | Not tested | Yes |
| **H3.3.1** | 7-11 | IF, RT-PCR | Yes | Tomov et al. 2016 | Tomov et al. | Tomov et al. | Yes |
| H3.1.1 | 8-11 | IF, RT-PCR | Yes | Tomov et al. 2016 | Tomov et al. | Tomov et al. | Yes |
| H3.4.1 | 6-9 | IF, RT-PCR | Not tested | This study | Not tested | Not tested | Yes |

| | | | | | | | |
|---|---|---|---|---|---|---|---|
| **A2.1.1** | 11-14 | IF, RT-PCR | Yes | Tomov et al. 2016 | Tomov et al. | Tomov et al. | Yes |
| A2.2.1 | 6-9 | IF, RT-PCR | Yes | This study | Not tested | Not tested | Yes |
| A2.2.2 | 10-17 | IF, RT-PCR | Yes | Tomov et al. 2016 | Tomov et al. | Tomov et al. | Yes |

**Supplementary Table 2.**

**Optimizing EMLO Conditions to Promote or Inhibit Elongation**

| **Condition** | **Promote** | **Inhibit** | **Comment** |
|---|---|---|---|
| Low cell number (300-400) | | | Starting aggregate |
| High cell number (9-10,000) | | | Starting aggregate |
| CHIR/FGF pretreatment as single cells | | | Advantage: homogenous NMps |
| CHIR/FGF pretreatment as 2D colonies | | | Advantage: mesendoderm, early NMp |
| CHIR/FGF longer induction | | | 3 days |
| CHIR/FGF shorter induction | | | 2 days |
| Direct to shaking cultures (day 0) | | | Advantage: EMLOs |
| Direct to 96-well plate static | | | Advantage: neuromuscular trunk organoids |
| Higher culture density | | | 6 well plate after day 5 |
| Lower culture density | | | 100 mm dish after day 5 |
| Fast shaking | | | 90 rpm |
| Slow shaking | | | 70 rpm |
| Early high-dose retinoic acid | | | Spinal cord neurospheres |
| Aggregate formation with spinal cord NSCs | | | Spinal cord neurospheres |

| | | | |
|---|---|---|---|
| Aggregate formation with anterior NSCs | | | Cerebral organoids |
| Dual SMAD Inhibition | | | Day 2 early addition |

**Supplementary Table 3. EMLO Qualitative Biomarker Summary and Signficance***

| Target | Neural (SC) | Mesoderm-Endoderm (ME, total) | Gut Tube Epithelium | Ganglion (Yes/No) | Significance | Distribution/ Comments |
|---|---|---|---|---|---|---|
| SOX2 | ++++ | ++ | + | No | Pluripotency, neural stem cells, gastrointestinal | Highly expressed in SC, basal levels in gut tube |
| CDH2 (N-Cadherin) | ++++ | + | - | Yes | Neural cell adhesion | SC rosettes and base of gut tube (anterior) |
| TUJ1 | ++++ | +++ | - | Yes | Post-mitotic neurons | SC, concentrated in EMT transition zone, gut tube envelopment, peripheral ganglia, other ME projections |
| GAP-43 | + | ++++ | - | Yes | Axon pathfinding | Axons enriched in ME |
| SMI312 | ++ | ++ | - | Yes | Axons | SC, ME |
| LHX9 | ++ | - | - | No | Dorsal spinal cord (dI1) | SC |
| LBX1 | ++ | - | - | No | Dorsal-intermediate spinal cord (dI4, dI6) | SC |
| PAX2 | ++++ | - | - | No | Dorsal-intermediate spinal cord (dI4, dI6, V0, V1) | SC |
| CHX10 | + | - | - | No | Ventral spinal cord interneuron (V2a) | SC |
| Nkx-6.1 | + | - | - | No | Ventral spinal cord motor neuron (MN) | SC |
| OPRM1 | +++ | ++ | + | No | Mu opioid receptor | SC apical aspect of rosettes, neurons at rosette base, gut tube lumen |
| GATA6 | - | ++++ | ++ | No | Heart fields, pancreas, diaphragm, liver, vasculature | ME gut tube plus mesenchyme |

| | | | | | | |
|---|---|---|---|---|---|---|
| GATA4 | - | ++++ | - | No | Endoderm specification, splanchnic mesoderm, valves, heart fields, diaphragm | ME mesenchyme only |
| FOXF1 | +++ | +++ | + | No | Splanchnic mesoderm, mesenchyme | SC, ME basal levels in gut tube, highly expressed in mesenchyme |
| ISL1 | +++ | ++ | + | Yes | MNs, dorsal INs, cardiac NCC, NCC-derived neurons | SC, gut tube, enriched at AIP and buds, peripheral ganglia |
| TFAP2α | ++++ | ++ | - | Yes | GABAergic neuronal progenitors, NCC, PNS | SC at base of rosettes, ME distribution along gut tube, peripheral ganglia, ME non-ganglia |
| SOX10 | ++++ | ++ | - | No | NCC, PNS | SC at base of rosettes, concentrated EMT transition zone, ME along gut tube |
| PAX7 | +++ | - | - | No | Dorsal spinal cord, myogenic progenitors | SC before EMT transition zone |
| PAX3 | + | ++ | - | No | Migratory NCC, myogenic progenitors | ME along gut tube; uniform in spheroids/ovoids |
| Vimentin | ++ | ++ | - | No | EMT, myogenic progenitors | EMT transition zone, ME |
| Peripherin | - | + | - | Yes | PNS (vs. CNS) | ME not SC |
| ChAT | + | + | - | Yes | Enteric NS, autonomic NS, motor neurons | Ganglion |
| Type IV Collagen | + | + | - | No | Basement membrane, EMT | EMT transition zone, basal lamina, gut tube, surface layer ME |
| FOXA2/ HNF-3β | + | +++ | +++ | No | Definitive endoderm | Spans SC to ME in gut tube |
| E-Cadherin | ++ | ++ | +++ | No | Epithelial | Gut tube cell-cell adhesion, surface layer ME |
| SOX17 | - | + | + | No | Definitive endoderm | Towards EMLO surface, migrating cells, gut tube |

| | | | | | | |
|---|---|---|---|---|---|---|
| Cytokeratin-19 | + | + | +++ | No | Definitive endoderm, surface ectoderm | Gut tube and surface layer |
| HAND1 | - | +++ | - | No | First heart field | ME mesenchyme |
| MEF2C | - | +++ | - | No | Second heart field | ME mesenchyme |
| Desmin | + | ++ | - | No | Myogenic progenitors | SC, ME |
| Troponin-T | - | + | - | No | Cardiomyocytes | Base of anterior intestinal portal-like region |

*Qualitative scale (++++, +++, ++, +, -) by day 22

#### SUPPLEMENTARY METHODS

**Neuromuscular organoids, cerebral organoids, neurospheres and small molecule manipulation.** The protocol for previously reported polarized neuromuscular trunk organoids (Faustino Martins et al., 2020) was validated using ED-hiPSC lines (**Supplementary Fig. 1**). Briefly, hiPSC colonies were dissociated with Accutase and seeded onto Matrigel as single cells at 100,000 /cm$^2$ in N2B27 supplemented with 3 µM CHIR, 40 ng/ml FGF2. Medium was changed every day for three days. At day -3, uniform NMps were dissociated with Accutase and forced to aggregate in 96 well plates (9-10,000 cells/well). At the time of dissociation, culture medium was changed to N2B27 supplemented with 10 ng/ml FGF2, 2 ng/ml HGF, 2 ng/ml IGF-1 (100 µl/well). 50 µM ROCK inhibitor was used at time of aggregation. One-half volume of medium was replaced with 100 µl N2B27 supplemented with 2 ng/ml HGF and IGF-1 24 h post-aggregation. After day 4, aggregates were maintained in N2B27 only. Day 10 organoids were transferred to a 60 mm dish and moved to an orbital shaker (70 rpm).

To test the effect of RA on EMLO elongation, 1 µM RA was added to shaking culture medium on day 2 of EMLO formation and maintained to day 13. In a separate experiment, we converted 2D adherent cultures that were pretreated for three days with 3 µM CHIR 99021 and 40 ng/ml FGF2 to spinal cord neural stem cells (scNSCs) by addition of 100 nM RA and 200 nM of the potent Hedgehog agonist Hh-Ag1.5. After 10 days, neurospheres were generated from adherent scNSCs on the orbital shaker at 75 rpm and were monitored for elongation.

Cerebral organoids were formed using a modified protocol from Trujillo et al. (2019) combined with the STEMCELL Technologies STEMdiff Cerebral Organoid and Maturation kit. Briefly, ~70% confluent hiPSC colonies maintained in mTeSR Plus were dissociated with Accutase diluted 1:1 in HBSS and transferred to 6-well plates treated with Anti-Adherence Rinsing Solution (STEMCELL Technologies) at ~4 x 10$^6$ cells per well. Cells were kept shaking at 95 rpm in mTeSR Plus supplemented with 1 µM Dorsomorphin (Tocris Bioscience) and 10 µM SB 431542 (Tocris Bioscience). 5 µM ROCK inhibitor was initially used. At day 3, medium was changed to N2B27 with the same small molecule and maintained for 7 days to achieve neural induction. We then switched neural expansion medium (STEMCELL Technologies) for 7 additional days, followed by maturation medium (STEMCELL Technologies) and maintained to day 50.

#### KEY RESOURCES TABLE

| REAGENT or RESOURCE | SOURCE | IDENTIFIER |
|---|---|---|
| **Antibodies** | | |
| Rat Anti-SSEA3 (AF488 conjugate) | STEMCELL Technologies | Cat.No.: 60061AD; RRID: AB_1118554 |
| Mouse anti-SSEA4 | STEMCELL Technologies | Cat.No.: 60062; RRID: AB_2721031 |
| Mouse anti-SOX2 | R&D Systems | Cat.No.: MAB2018; RRID: AB_358009 |
| Goat anti-SOX2 | R&D Systems | Cat.No.: AF2018; RRID: AB_355110 |
| Goat anti-Brachyury | R&D Systems | Cat.No.: AF2085; RRID: AB_2200235 |
| Goat anti-GATA4 | R&D Systems | Cat.No.: AF2606; RRID: AB_2232177 |
| Goat anti-GATA6 | R&D Systems | Cat.No.: AF1700; RRID: AB_2108901 |
| Mouse anti-CDX2 | DSHB | Cat.No.: PCRP-CDX2-1A3; RRID: CVCL_KD04 |
| Rabbit anti-ISL1 | Sigma-Aldrich | Cat.No.: HPA057416; RRID: AB_2683431 |
| Mouse anti-TFAP2a | DSHB | Cat.No.: 3B5; RRID: AB_667767 |
| Goat anti-SOX10 | R&D Systems | Cat.No.: AF2864; RRID: AB_442208 |
| Mouse anti-Collagen IV $\alpha$1 | R&D Systems | Cat.No.: MAB6308 |
| Rabbit anti-Collagen I $\alpha$1 | Novus Biologicals | Cat.No.: NBP1-30054; RRID: AB_1968486 |
| Goat anti-Vimentin | R&D Systems | Cat.No.: AF2105; RRID: AB_355153 |
| Goat anti-CDH1/E-cadherin | R&D Systems | Cat. No.: AF749; RRID: AB_355568 |
| Goat anti-FOXA2 | R&D Systems | Cat.No.: AF2400; RRID: AB_2294104 |
| Rabbit anti-FOXF1 | Abcam | Cat.No.: ab168383 |
| Rabbit anti-EOMES | Abcam | Cat.No.: ab23345; RRID: AB_778267 |
| Mouse anti-Cytokeratin 19 | R&D Systems | Cat.No.: MAB3506 |
| Goat anti-SOX17 | R&D Systems | Cat.No.: AF1924; RRID: AB_355060 |
| Rabbit anti-$\beta$-tubulin III/TUJ1 | BioLegend | Cat.No.: 802001; RRID: AB_2564645 |
| Mouse SMI312 cocktail | BioLegend | Cat.No.: 837904; RRID: AB_2566782 |
| Mouse anti-GAP-43 | Encor | Cat.No.: MCA-5E8; RRID: AB_2572287 |

| | | |
|---|---|---|
| Mouse anti-Peripherin | Santa Cruz Biotechnologies | Cat.No.: sc-377093 |
| Mouse anti-ChAT | R&D Systems | Cat.No.: MAB3447 |
| Rabbit anti-CDH2/N-cadherin | Cell Signaling Technologies | Cat.No.: 13116; RRID: AB_2687616 |
| Mouse anti-ZO1 | Invitrogen | Cat.No.: 33-9100; RRID: AB_2533147 |
| Rabbit anti-Mu Opioid Receptor (OPRM1) | Abcam | Cat.No.: ab10275; RRID: AB_2156356 |
| Rabbit anti-GAD65&67 | Abcam | Cat.No.: ab11070; RRID: AB_297722 |
| Rabbit anti-TLX3 | Abcam | Cat.No.: ab184011 |
| Rabbit anti-LBX1 | Invitrogen | Cat.No.: PA564884; RRID: AB_2662958 |
| Rabbit anti-PAX2 | Invitrogen | Cat.No.: 716000; RRID: AB_2533990 |
| Goat anti-PAX3 | R&D Systems | Cat.No.: AF2457; RRID: AB_416599 |
| Mouse anti-PAX7 | R&D Systems | Cat.No.: MAB1675; RRID: AB_2159833 |
| Rabbit anti-LHX9 | Abcam | Cat.No.: ab224357 |
| Mouse anti-CHX10 | Santa Cruz Biotechnologies | Cat.No.: sc-374151 |
| Mouse anti-NKX6.1 | DSHB | Cat.No.: F55A12; RRID: AB_532379 |
| Rabbit anti-BRN3A | EMD Millipore | Cat.No.: AB5945; RRID: 92154 |
| Goat anti-BMP2/4 | R&D Systems | Cat.No.: AF355 |
| Mouse anti-MEF2C | R&D Systems | Cat.No.: MAB6786 |
| Goat anti-HAND1 | R&D Systems | Cat.No.: AF3168 |
| Goat anti-Desmin | R&D Systems | Cat.No.: AF3844; RRID: AB_2092419 |
| Mouse anti-Troponin T (cardiac) | R&D Systems | Cat.No.: MAB1874 |
| **Chemicals** | | |
| mTeSR Plus | STEMCELL Technologies | Cat.No.: 05825 |
| mFreSR | STEMCELL Technologies | Cat.No.: 05854 |
| DMEM/F-12 | Thermo Fisher Scientific | Cat.No.: 11320033 |
| Knockout DMEM | Thermo Fisher Scientific | Cat.No.: 10829018 |
| Knockout Serum Replacement | Thermo Fisher Scientific | Cat.No.: 10828028 |
| Neurobasal Plus Medium | Thermo Fisher Scientific | Cat.No.: A3582901 |

| | | |
|---|---|---|
| BrainPhys Neuronal Culture Medium | STEMCELL Technologies | Cat.No.: 05790 |
| N-2 supplement (100X) | Thermo Fisher Scientific | Cat.No.: 17502048 |
| B-27 supplement (50X) | Thermo Fisher Scientific | Cat.No.: 17504044 |
| GlutaMAX | Thermo Fisher Scientific | Cat.No.: 35050061 |
| MEM Non-Essential Amino Acids | Thermo Fisher Scientific | Cat.No.: 11140050 |
| Penicillin-Streptomycin | Thermo Fisher Scientific | Cat.No.: 15140122 |
| hESC-qualified Matrigel | Corning | Cat.No.: 08-774-552 |
| CHIR 99021 | Tocris Bioscience | Cat.No.: 4423 |
| bFGF/FGF2 | R&D Systems | Cat.No.: 233-FB |
| FGF8 | R&D Systems | Cat.No.: 423-F8 |
| HGF | R&D Systems | Cat.No.: 294-HG |
| IGF-1 | R&D Systems | Cat.No.: 291-G1 |
| BDNF | R&D Systems | Cat.No.: 248-BDB-005 |
| GDNF | R&D Systems | Cat.No.: 212-GD-010 |
| Dibutyryl-cyclic AMP | Sigma-Aldrich | Cat.No.: D0260 |
| Y-27632 | Tocris Bioscience | Cat.No.: 1254 |
| Retinoic acid | Sigma-Aldrich | Cat.No.: R2625 |
| Hh-Ag1.5 | Cellagen Technology | Cat.No.: C4412-2s |
| LDN 193189 | Tocris Bioscience | Cat.No.: 6053 |
| SB 431542 | Tocris Bioscience | Cat.No.: 1614 |
| DAPT | Tocris Bioscience | Cat.No.: 2634 |
| SU 5402 | Tocris Bioscience | Cat.No.: 3300 |
| Dorsomorphin | Tocris Bioscience | Cat.No.: 3093 |
| Accutase | STEMCELL Technologies | Cat.No.: 07920 |
| Gentle Cell Dissociation Reagent | STEMCELL Technologies | Cat.No.: 07174 |
| Neural Tissue Dissociation Kit (P) | Miltenyi Biotec | Cat.No.: 130-092-628 |
| Anti-Adherence Rinsing Solution | STEMCELL Technologies | Cat.No.: 07010 |
| Fluo-4 AM | Thermo Fisher Scientific | Cat.No.: F14201 |

| | | |
|---|---|---|
| HBSS | Thermo Fisher Scientific | Cat.No.: 14025076 |
| HBSS CM-free | Thermo Fisher Scientific | Cat.No.: 14175079 |
| DAMGO mu opioid receptor agonist | Abcam | Cat.No.: ab120674 |
| Naloxone hydrochloride | Abcam | Cat.No.: ab120074 |
| Histodenz | Sigma-Aldrich | Cat.No.: D2158 |
| Bovine Serum Albumin Fraction V | Fisher Scientific | Cat.No.: BP1600-100 |
| Triton X-100 | Electron Microscopy Sciences | Cat.No.: 221440 |
| NucBlue fixed cell stain | Thermo Fisher Scientific | Cat.No.: R37606 |
| **Commercial Assays** | | |
| STEMdiff Cerebral Organoid and Maturation Kit | STEMCELL Technologies | Cat.No.: 08570-1 |
| **Experimental Models: Cell Lines** | | |
| hiPSC lines used in this study | Paluh lab | Supplementary Table 1 |
| **Software** | | |
| Keynote | Apple | https://www.apple.com/keynote/ |
| GraphPad Prism 9 | GraphPad | https://www.graphpad.com/scientific-software/prism/ |
| Excel | Microsoft | https://www.microsoft.com/en-us/microsoft-365/excel |
| Fiji | Schindelin et al., 2012 | https://imagej.net/Fiji |
| Illustrator CC2020 | Adobe | https://www.adobe.com/products/illustrator.html |
| Imaris | BitPlane | https://imaris.oxinst.com/ |
| cellSens | Olympus | https://www.olympus-lifescience.com/en/software/cellsens/ |
| **Other** | | |
| 6 well plate | CELLTREAT | Cat.No.: 229105 |
| 100 mm petri dish | Fisher | Cat.No.: S33580A |
| Lab-Tek II chambered coverglass | Thermo Fisher Scientific | Cat.No.: 155382 |
| CHEMcell VERSA-ORB2 orbital shaker | Chemglass Life Sciences | Cat.No.: CLS-4021-100 |

### Supplementary Figure 1

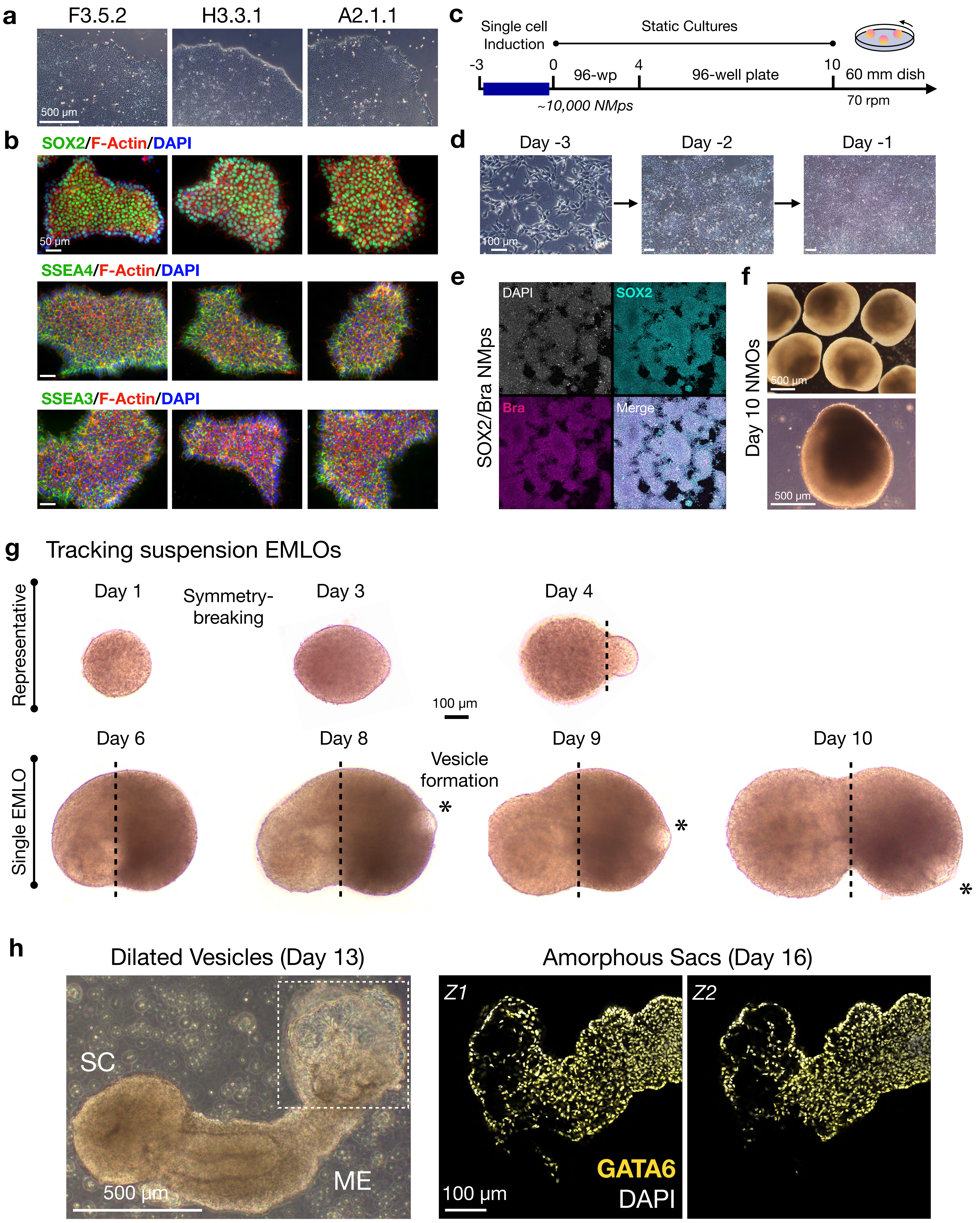

### Supplementary Figure 2

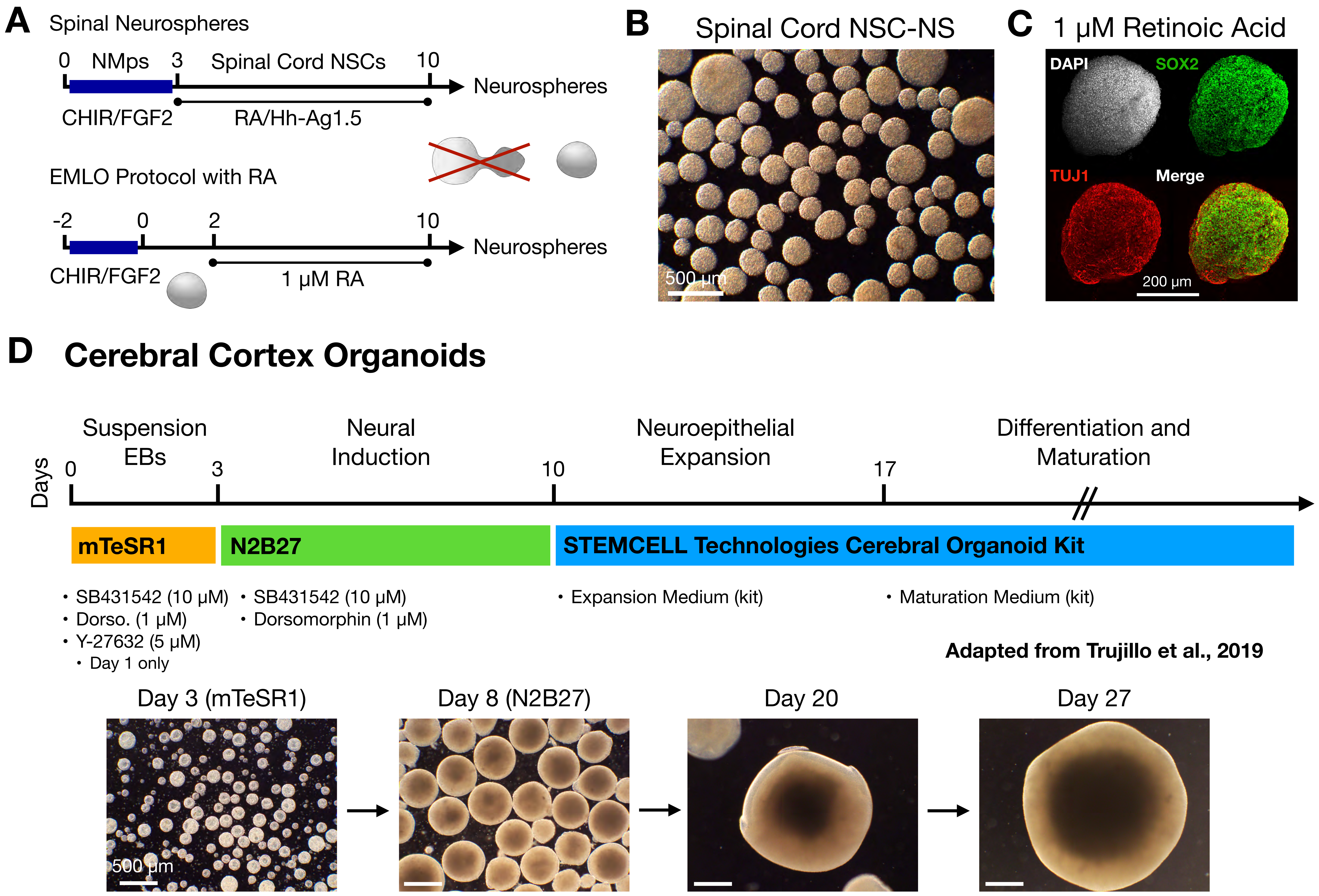

**Supplementary Figure 3**

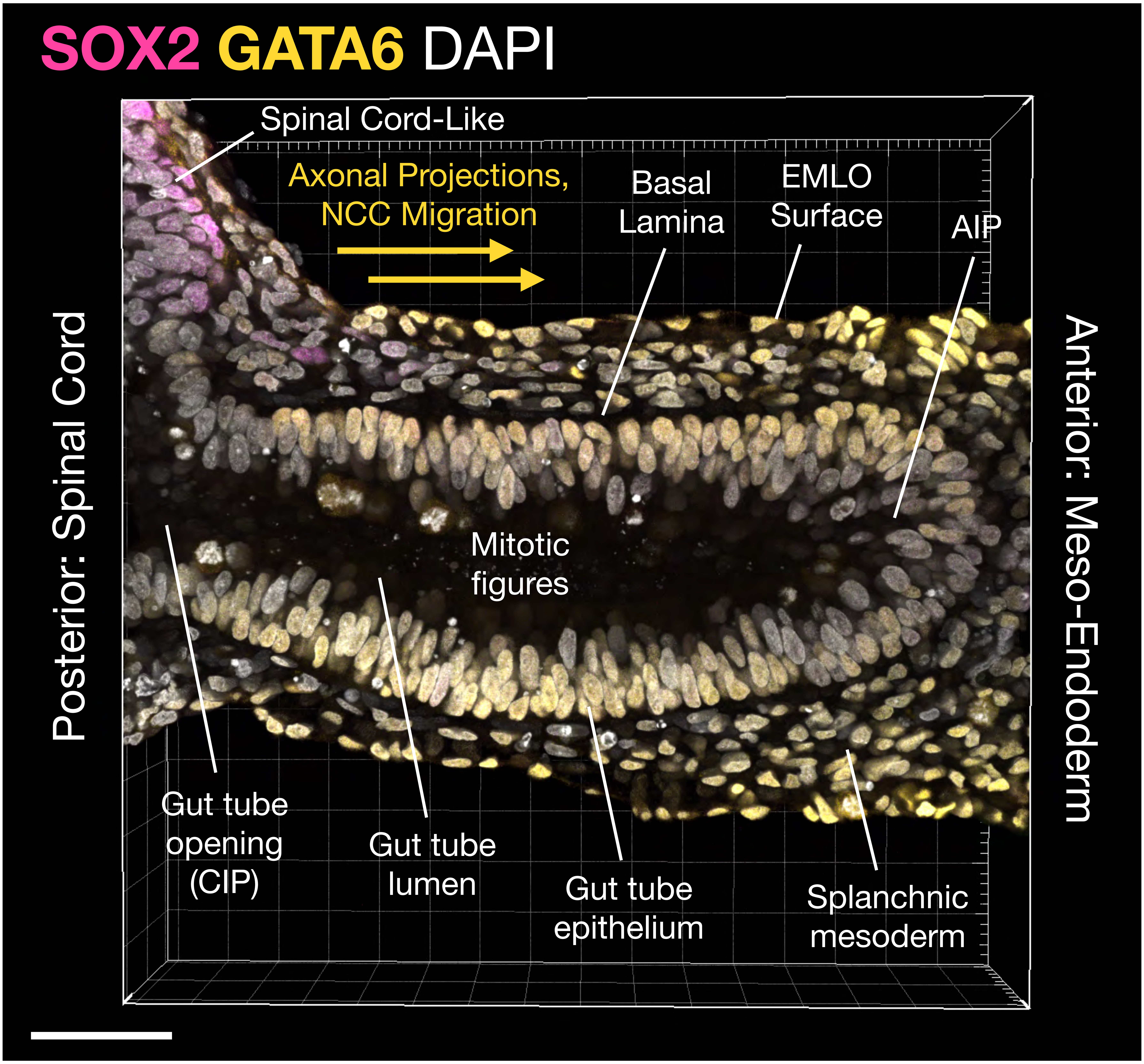

#### Supplementary Figure 4

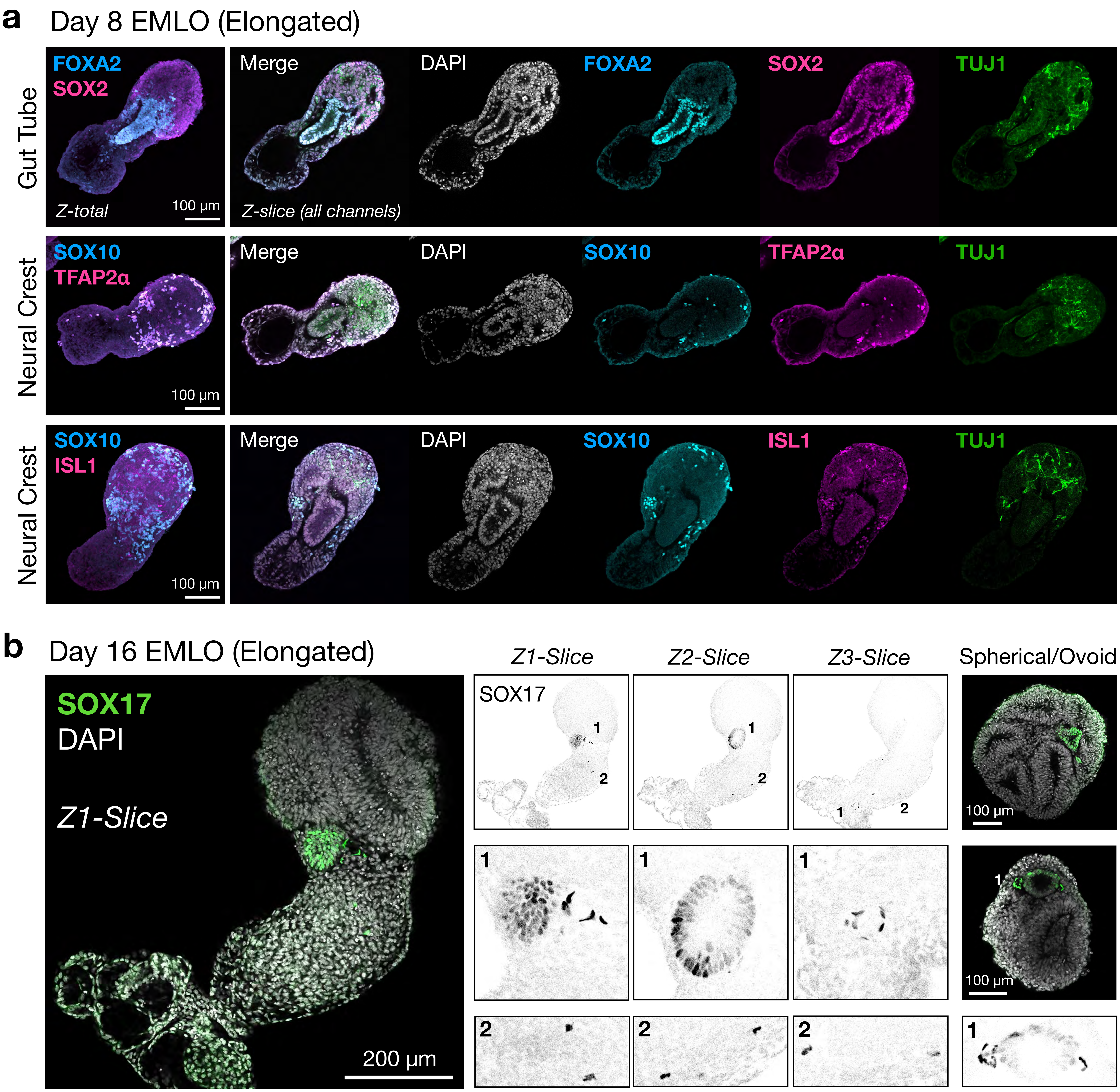

### Supplementary Fig. 5

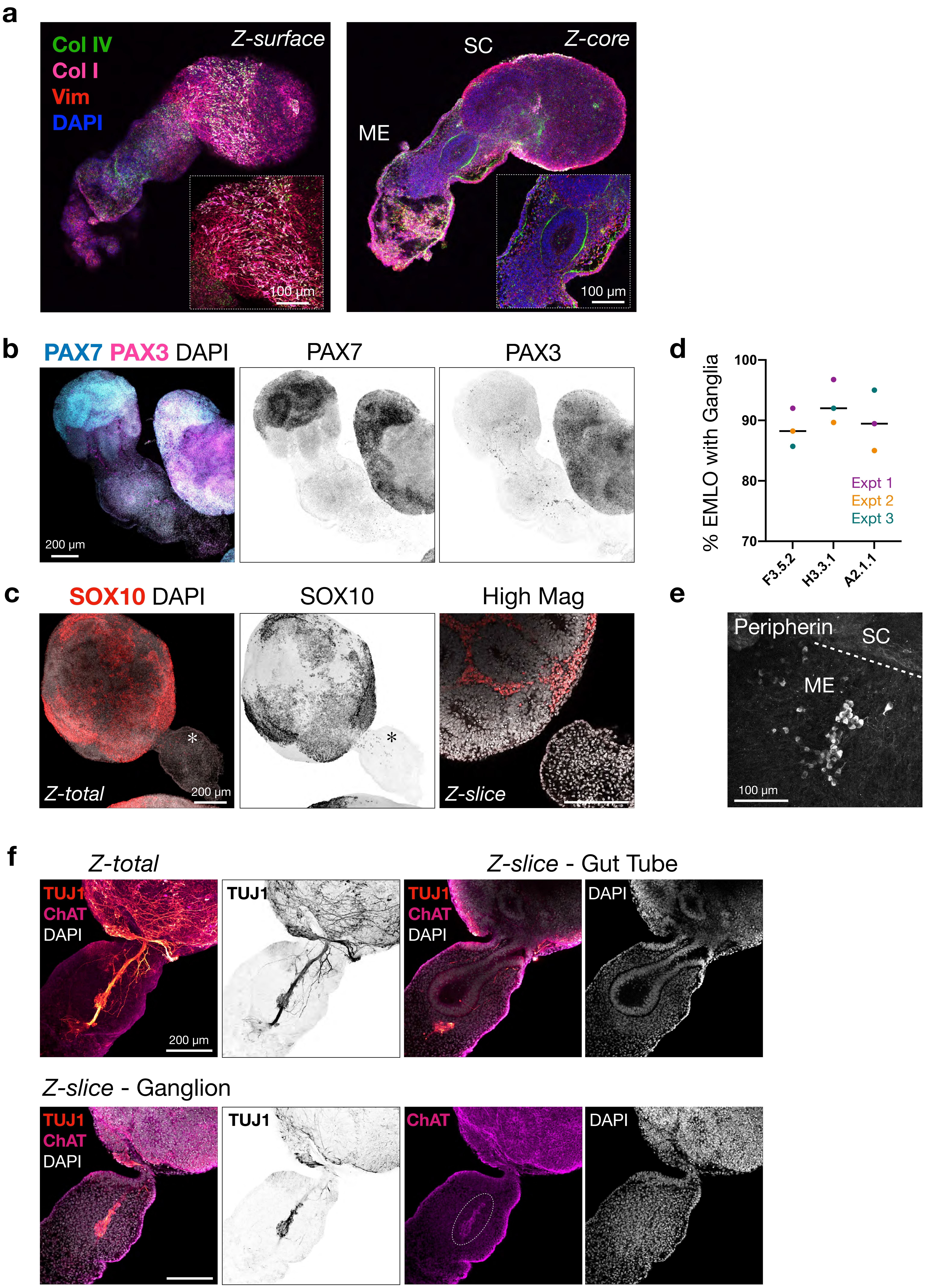
